# Dust storm-driven dispersal of potential pathogens and antibiotic resistance genes in the Eastern Mediterranean

**DOI:** 10.1101/2024.06.24.600361

**Authors:** Burak Adnan Erkorkmaz, David Zeevi, Yinon Rudich

**Author notes:** Corresponding Author: Yinon Rudich.

## Abstract

The atmosphere hosts a microbiome that connects distant ecosystems yet remains relatively unexplored. In this study, we tested the hypothesis that dust storms enhance the spread of pathogenic microorganisms and whether these microorganisms carry antibiotic resistance and virulence-related genes in the Eastern Mediterranean. We collected air samples during a seasonal transition period, capturing data from 13 dusty days originating from Middle Eastern sources and 32 clear days, with temperatures ranging from 16.5 to 27.1 °C. Using metagenomic analysis, we identified several facultative pathogens like *Klebsiella pneumoniae*, *Stenotrophomonas maltophilia*, and *Aspergillus fumigatus*, which are linked to human respiratory diseases, and others like *Zymoseptoria tritici*, *Fusarium poae*, and *Puccinia striiformis*, which are harmful to wheat. The abundance of these pathogens increased during dust storms and with rising temperatures. Although we did not find strong evidence that these species harbored antibiotic resistance or virulence-related genes, which could be linked to their pathogenic potential, dust storms transported up to 125 times more total antibiotic resistance genes, as measured by RPKM abundance, compared to clear conditions. These levels during dust storms far exceeded those found in other ecosystems. While further research is needed to determine whether dust storms and temperature variations pose an immediate threat to public health and the environment, our findings underscore the importance of continuous monitoring of atmospheric microbiomes. This surveillance is crucial for assessing potential risks to human health and ecosystem stability, particularly in the face of accelerating global climate change.

## Introduction

The atmospheric microbiome plays a vital role in Earth’s ecosystems, contributing to important processes such as ice nucleation and atmospheric chemical reactions ^1–6^. Despite its significance, the atmospheric microbiome is less studied compared to other habitats, particularly concerning its ecological and health-related impacts. The atmospheric microbiome is remarkably diverse and dynamic, containing microorganisms from multiple sources. Some microbes are naturally released into the air through processes like wind and oceanic aerosolization, while others come from human activities, such as industrial operations, agriculture, and transportation ^7–9^. Dust storms act as carriers, transporting large amounts of particles from arid regions and dispersing microorganisms across vast distances ^10,11^. This process not only alters the composition and dynamics of the atmospheric microbiome but also connects distant ecosystems ^12–14^. Dust storms introduce a wide variety of soil microorganisms, significantly increasing atmospheric microbial abundance during these events ^12,13,15^. Once the dust settles, it can affect both terrestrial and aquatic ecosystems ^16,17^. The presence of potentially harmful microorganisms in dust raises concerns about risks to public health and ecosystems ^12,13,18^. Previous studies have shown that pathogens can be transported over long distances through the atmosphere, contributing to plant diseases ^19–21^. Additionally, epidemiological research has linked dust storms to human health issues, particularly respiratory diseases ^22–28^. This raises the question of whether biological agents within dust are responsible for triggering these diseases, an aspect of the atmospheric microbiome that remains underexplored. Although antibiotic resistance is typically associated with hospitals due to prolonged medication use, genes related to antibiotic resistance have also been detected in airborne dust ^12,13^. This suggests that the atmosphere may serve as a vehicle for spreading genes across distant regions. Thus, it is essential to investigate whether dust storms carry potential pathogens or genes linked to antibiotic resistance and virulence. This is especially important as climate change is expected to increase particulate matter and lead to more frequent dust events, particularly in the Eastern Mediterranean ^29,30^. The poleward expansion of Hadley cells is also predicted to worsen drying conditions and extend dust storms further into Europe ^29^. Understanding these dynamics is crucial for assessing the potential impact of climate change on the atmospheric microbiome at local and global scales.

Studying the atmospheric microbiome presents unique challenges due to its low-biomass nature ^31^. Overcoming this challenge requires sophisticated bioinformatic algorithms and high sequencing depth for accurate analysis. While traditional methods like amplicon sequencing, which target specific genetic markers such as the 16S ribosomal RNA gene and internal transcribed spacer (ITS), have been utilized to identify taxonomic features, they typically provide resolution only at the genus level for bacteria and fungi ^12,13,15,32^. Recent studies have employed shotgun metagenomic and metatranscriptomic sequencing in other ecologies to explore the atmospheric microbiome, utilizing advanced aerosol sampling devices and genomic technologies ^31,33–35^. However, most research on the dust microbiome has focused on shallow taxonomic characterization or generalized functional analysis ^12,13,15,32,36,37^, rather than tailoring bioinformatic tools that are specific to each scientific research question. Moreover, some studies have quantified fragments of a few known antibiotic resistance genes by qPCR, overlooking the broader diversity, richness, and origin of these genetic features ^12,13^. Furthermore, the sampling strategies in these studies have often lacked longitudinal designs, which are necessary to capture the dynamic relationship between atmospheric conditions and microbial features over time. Therefore, we need comprehensive research to understand how dust storms and temperature fluctuations specifically influence the dispersal of pathogens and whether these pathogens carry traits related to pathogenicity.

In this study, we collected air samples during a seasonal transition period, capturing data from 13 dusty days from prominent Middle Eastern dust sources and 32 clear atmospheric conditions, with temperatures ranging from 16.5 to 27.1 °C. Our primary objective is to use metagenomic approaches to identify potential pathogens in these atmospheric samples and assess whether there is a link between these species, dust storms, and rising temperatures. Additionally, we aim to determine if these pathogens carry genes associated with antibiotic resistance and virulence, which could present risks to both public health and the environment.

## Materials and Methods

### Sample collection

We collected a total of 45 air samples on the rooftop of a four-story building at the Weizmann Institute of Science in Rehovot, Israel (31.9070 N, 34.8102 E; 80 m above mean sea level). This timeframe allowed us to capture the diverse dynamics of air masses affected by distinct Eastern and Western dust storm origins, as well as the fluctuations in temperature ranging from 16.5 to 27.1 °C. We predicted dust storms using various online forecast tools and collected meteorological data, dust column density maps, and air mass back trajectories to estimate the origins of dust and air masses. For metagenomics analysis, we employed the Coriolis µ microbial air sampler (Bertin Technologies) to collect air samples for 2 hours, at a flow rate of 200 L/min. Detailed information on atmospheric data collection, air sampling (including control samples), the decontamination procedures applied, as well as results of the shotgun metagenomic analysis of control samples, can be found in the Supporting Information.

### DNA Extraction

We extracted DNA from the filters using a PowerWater DNA isolation kit (Qiagen, Dresden, Germany) following the manufacturer’s protocol. Following this, we quantified the extracted DNA from the samples using a Qubit 2.0 fluorometer, employing the Qubit dsDNA HS (High Sensitivity) Assay Kit (Invitrogen) with dsDNA HS buffer. Detailed information regarding the DNA yield obtained from the control samples can be found in the Supporting Information.

### DNA Library Preparation and Metagenomic Sequencing

Shotgun metagenome libraries were generated using a Nextera XT library preparation kit according to the manufacturer’s instructions (Illumina). The libraries were pooled and sequenced using high-output flow cells with paired-end 2 × 150 base reads on an Illumina NovaSeq6000 sequencer, utilizing an S4 flow cell. The library preparation and sequencing processes were conducted at the Rush Genomics and Microbiome Core Facility (GMCF) located at Rush University.

### Data processing and metagenomic analysis

We performed a comprehensive quality assessment of the metagenomic datasets using FASTQC software (version 0.11.9, www.bioinformatics.babraham.ac.uk/projects/fastqc/). We applied quality trimming and Illumina adapter removal using Trimmomatic ^38^ with specific parameters: ILLUMINACLIP:TruSeq3-PE.fa:2:30:10 LEADING:20, TRAILING:20, SLIDINGWINDOW:4:15, and MINLEN:35. The application of Trimmomatic’s quality control steps resulted in an average of 4.3 million high-quality reads per sample (3-11 million paired-end) that met the predefined quality criteria, as presented in Supporting Tables 1 and 2.

We performed taxonomic classification of the metagenomic data using Kraken2 (v2.1.2) with exact k-mer matching ^39^. We further refined these results using Bracken (v2.7.0) ^40^, and we evaluated the Kraken2 results by estimating the number of distinct k-mers associated with each taxon using KrakenUniq ^41^. Details of these analyses can be found in the Supporting Information. To further verify the specificity of taxonomic classification result, we conducted read mapping from each sample to reference genomes, focusing particularly on low-abundance species such as potential pathogens detected in air samples. We obtained the reference genomes of the thirty-four most prevalent potential pathogens, and generally the most abundant taxa across all samples, with *G. obscurus* as a reference, from the NCBI RefSeq database on February 12, 2024. Subsequently, we aligned all reads from the samples to each reference genome separately using Bowtie2 (--very-sensitive-local) ^42^. The mapping results of each individual sample were sorted and converted to BAM format using SAMtools ^43^. We reported genome coverage for all positions and the fractions of genome covered by reads using the bedtools genomecov command (-bga) and (-max 50), respectively ^44^. Finally, we visualized genome tracks using the ‘circlize’ package in R ^45^.

We performed functional characterization of the metagenomics data using the BLASTX function in DIAMOND (v2.1.7.161) ^46^, utilizing merged paired-end reads and specific parameters (--very-sensitive -- id 80 --query-cover 80) to enhance sensitivity. For this analysis, we employed the NCBI-nr database from June 2023. Subsequently, we processed the DIAMOND results further using the daa-meganizer tool (-- minScore 80 --minPercentIdentity 80 --minSupport 5) in MEGAN (version 6.24.20) ^47^. We assigned functional features when at least five reads uniquely aligned with a specific feature. To annotate the NCBI-nr data into SEED categories, we utilized the megan-map-Feb2022.db file.

We conducted metagenomic assembly using metaSPAdes (version 3.15.3) ^48^ on quality-trimmed, paired-end reads, employing default settings. Prediction of open reading frames (ORFs) for protein-coding genes from assembled contigs was achieved using Prodigal (version 2.6.3) ^49^ with the ‘-p meta’ option, enabling anonymous mode as recommended for metagenomic datasets.

We carried out antibiotic resistance and virulence profiling of the metagenomic data using hmmscan (HMMER, version 3.3.2-gompic-2020b) ^50^ with a stringent e-value threshold of 1×10^−5^ to predict the function of ORFs. We performed separate searches for genes containing Pfam modules ^51^ related to ‘antibiotic resistance’, ‘virulence’, and ‘pathogenicity’. We retained only the top matches in the hmmscan results when the same ORFs yielded multiple hits to the same Pfam entry or the same contig had more than one hit, which was rare due to short contig lengths.

To estimate the abundance of all contigs, including those associated with antibiotic resistance and virulence, we mapped the quality-trimmed, paired-end reads to the contigs in each sample, providing coverage information. This mapping was performed using bbmap.sh (BBMap, v.38.90) with default settings. Subsequently, we utilized pileup.sh to convert the coverage data into reads per kilobase per million mapped reads (RPKM). This normalization procedure ensures accurate rescaling to correct for both library size and gene length. We then conducted taxonomic annotation on these contigs using Kraken2 with default parameters, employing the same custom database constructed for Kraken2. We obtained the complete lineage of taxonomy for each contig from the kingdom level to the species level using Taxonkit with the taxdump file downloaded from the NCBI database on February 27, 2024 ^52^. We created the Sankey diagrams for visualization using the ‘networkD3’ package in R.

To address variations in library size across samples, we normalized the read counts obtained from tools such as Bracken and DIAMOND to counts per million (CPM) values. We performed this normalization by calculating CPM values, where each value represents the relative proportion of reads (i.e., reads normalized for library size through total sum scaling) multiplied by 1,000,000. Our metagenomics data deviated from the fundamental assumption that most features should exhibit minimal changes between samples. Consequently, we refrained from employing alternative inter-sampling normalization approaches. Instead, we opted for CPM normalization as it adequately accounts for library size differences and facilitates reliable comparisons of read abundances across samples. We then calculated the richness (observed number of features) and the Shannon-Wiener diversity index. Additionally, we conducted Principal Coordinates Analysis (PCoA) and permutational MANOVA using Bray-Curtis dissimilarity to analyze the taxonomic and functional features of the compositional data. Further details and the rationale behind the statistical analyses are provided in the Supporting Information.

## Data availability

The metagenomic sequences utilized in this study have been deposited in the National Center for Biotechnology Information (NCBI) database under the accession number PRJNA1045528. Additionally, Supporting Table 8, has been archived in a suitable repository, accessible at (https://doi.org/10.5281/zenodo.11090298).

## Results

### Abundance, richness, and diversity of the atmospheric microbiome

We observed variations in the total sampled biomass, as measured by the total DNA yield obtained from air samples. We found significantly higher concentrations in dusty atmospheric conditions (2.10±4.43 ng µL^-1^, mean±SD) compared to clear atmospheric conditions (0.31±0.33 ng µL^-1^) (Mann–Whitney *U* test, *p*<0.001, Supporting Figure 1). Due to the inherently low biomass nature of air samples, we did not obtain sufficient DNA for both genomic sequencing analysis and qPCR analysis. qPCR provides a more sensitive measure of biomass compared to fluorometric analysis as in Qubit. This limitation is particularly relevant when studying dust storms, as the sampling window to capture the storm at its peak is typically limited to a few hours. However, this finding is consistent with previous qPCR quantifications from our earlier studies, which employed longer sampling times. This extended sampling likely compromised the specificity of the microbial characteristics associated with dust storms. These earlier studies also demonstrated an increase in absolute microbial abundance on dusty days compared to clear conditions ^12,13,15^. We provide a detailed discussion of the DNA yield from operational blanks, meteorological conditions before and during the sampling period, and the estimated origin of dust in the Supporting Information, including Supporting Figures 2 and 3, and Supporting Table 3.

We found that only a small portion of the total reads identified with a known taxonomy (3.81±3.37%, mean±SD, Supporting Figure 4). Bacteria predominantly comprised the atmospheric microbial community, accounting for 95.8±1.94% (mean±SD) of the assigned reads, while fungi, archaea, other eukaryotes, and viruses constituted minor proportions across all samples (3.79±1.8%, 0.12±0.31%, and 0.12±0.22%, respectively, Supporting Figure 5). Notably, specific bacterial classes, including Actinomycetes (49.0±10.4%, mean±SD), Gammaproteobacteria (17.9±10.3%), and Alphaproteobacteria (15.5±3.57%), emerged as dominant taxa within the atmospheric microbial community (Supporting Figure 6).

We observed variations in the composition of prominent microbial classes under different atmospheric conditions (Supporting Figure 7). Dusty conditions showed significant increases in certain classes such as Actinomycetes (Mann–Whitney *U* test, *p*=0.068) and Rubrobacteria (*p*=0.006), known for their resilience to environmental stressors ^53^. Notably, *Geodermatophilus obscurus*, a prominent Actinomycetes species, exhibited distinct dominance in dusty conditions compared to clear ones (Mann–Whitney *U* test, *p*<0.001), constituting 16.7±11.2% (mean±SD) of the microbial community in dusty conditions and 4.97±3.85% in clear conditions (Supporting Figure 8). Conversely, certain Hydrobacteria group microorganisms, including Alphaproteobacteria (Mann–Whitney *U* test, *p*<0.001) and Betaproteobacteria (*p*=0.055), showed increased relative abundance in clear atmospheric conditions, suggesting a stronger marine influence ^53^.

We analyzed how various environmental factors contribute to the observed variability in the atmospheric microbiome using variance partitioning analysis (Adonis). Our results show that daily mean PM_10_ concentration explained the largest proportion of total variation (9.8% for taxonomy, 8.8% for functional attributes). This was followed by mean temperature and air mass origin (temperature: 9.3% and 4.6%; air mass: 9.3% for taxonomy, but not functional attributes, Supporting Table 4). Notably, relative humidity did not significantly influence this variation. Additionally, our PCo analysis revealed distinct microbial patterns. Firstly, air samples collected on clear days formed a separate cluster compared to those from dusty days, regardless of air mass origin (Adonis, *R*^2^=0.09, *p*<0.001). On dusty days, the microbiomes of southwesterly air masses partially overlapped with those from clear days, while samples from easterly air masses were distinct from both clear days and southwesterly air masses. Overall, microbiomes from clear days tended to cluster together, whereas those from dusty days were more dispersed and less cohesive. Second, we observed a temperature-related trend, particularly under clear atmospheric conditions. This was supported by a significant correlation between the PCo1 and temperature (Pearson’s correlation, *r*=0.44, *p*=0.002, Figure 1A and Supporting Figure 9A). Additionally, we observed similar, yet less pronounced trend in PCo analysis of the functional characteristics of the atmospheric microbiome (Figure 1D and Supporting Figure 9B). We also found significant differences in functional attributes between the microbiomes of clear and dusty conditions, particularly in the SEED sub-categories related to cell envelope (Adonis, *R*^2^=0.14, *p*<0.001) and metabolism (*R*^2^=0.09, *p*<0.001), and stress response, defense, and virulence (*R*^2^=0.12, *p*<0.001, Supporting Figure 10). However, the functional dissimilarity between atmospheric microbiomes was significantly lower than the taxonomic dissimilarity between samples (Mann–Whitney *U* test, *p*<0.001, Supporting Figure 11). This finding suggests that functional characteristics are more stable and resilient than taxonomic traits in the atmospheric microbiome.

**Figure 1.**
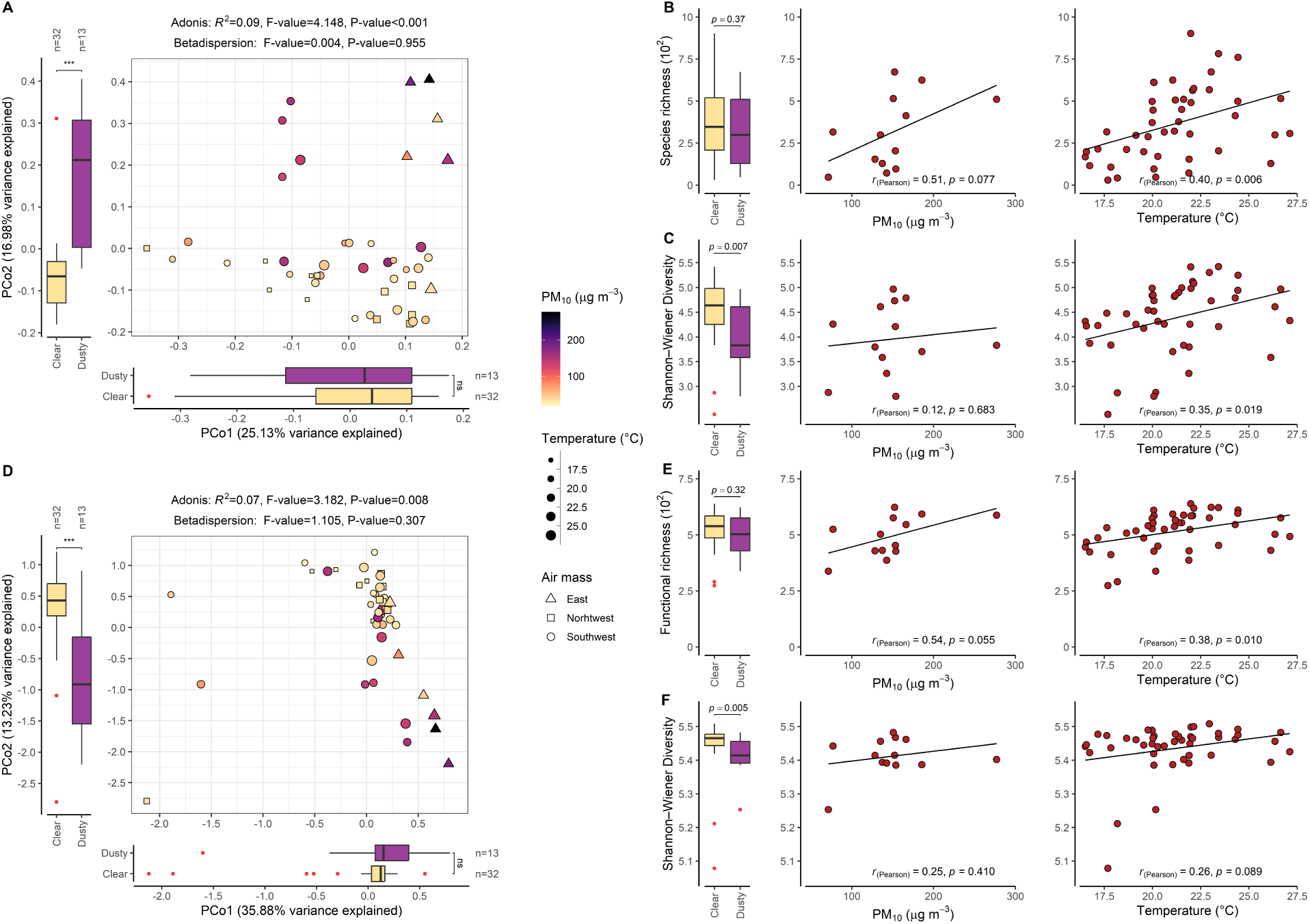
Taxonomic and functional characteristics of the atmospheric microbiome in relation to environmental variables. (A, D) Principal Coordinate Analysis (PCoA) plots illustrate Bray–Curtis dissimilarity of taxonomic (A) and functional (D) features, demonstrating the impact of environmental variables on compositional data. Axes indicate the variance proportion explained by respective PCoAs. Adonis test results, based on Bray–Curtis dissimilarity, are presented below each plot’s title. Multivariate homogeneity of group dispersions (betadispersion), a prerequisite of permutational MANOVA, is also displayed. Mann–Whitney *U* test assessed significant differences in PCoA axes, richness, and diversity between dusty and clear conditions. Significance levels are indicated by symbols: ‘***’ for <0.001, ‘**’ for <0.01, ‘*’ for <0.05, ‘.’ for <0.1, and ‘ns’ for <1. (B, E) Richness plots exhibit species (B) and functional (E) richness, depicting the observed number of features, while (C, F) diversity plots display species (C) and functional (F) diversity using the Shannon–Wiener diversity index for the atmospheric microbiome under dusty and clear conditions. Pearson’s correlation analysis explores the relationship between PM_10_ concentration (exclusive to dusty atmospheric conditions), temperature across all samples, and microbiome richness/diversity indices. Boxplots display median values (center lines), first and third quartiles (lower and upper hinges, 25^th^ and 75^th^ percentiles), while whiskers extend from hinges to values within 1.5 times the Interquartile Range (IQR). Outliers are red dots.

During the sampling campaign, daily mean temperatures on dusty days were generally higher (Mann– Whitney *U* test, *p*=0.008, Supporting Figure 12). Our analysis, including PCoA and Adonis, showed that PM_10_ concentration was the primary factor driving community composition. Moreover, during the major easterly dust storms on April 24-26, 2022, we observed unique microbial features in the atmospheric microbiome for the first time in this campaign. This event, lasting over three days, saw PM_10_ concentrations exceed 900 μg m^−3^. It is plausible that this extreme period affected local ecosystems, potentially contributing to the appearance of these new taxa in subsequent samples. We also found that temperature exerted a secondary, albeit lesser, influence, especially toward the end of the campaign. This overlap between PM_10_ concentration and temperature complicates our efforts to analyze their individual effects on the atmospheric microbiome. To address this, we assessed the impact of both PM_10_ and temperature variations across all samples, while also analyzing the specific effect of PM_10_ on dusty days separately.

We found that the richness of the atmospheric microbiome, in terms of taxonomic and functional attributes, was similar on both clear and dusty days (Mann–Whitney *U* test, *p*=0.37 and *p*=0.32, respectively, Figure 1B and 1E). However, during dusty conditions, we observed a trend of increasing richness correlated with higher PM_10_ concentrations, although this trend did not achieve statistical significance (Pearson’s correlation, *r*=0.51, *p*=0.077 and *r*=0.54, *p*=0.055). In contrast, clear days did not exhibit a similar trend (Pearson’s correlation, *r*=0.28, *p*=0.123 and *r*=0.18, *p*=0.312, respectively), likely due to the limited range of PM_10_ concentrations on those days. However, we did find that the richness of the atmospheric microbiome increased with rising temperatures across all samples (Pearson’s correlation, *r*=0.40, *p*=0.006 and *r*=0.38, *p*=0.010, respectively) and specifically on clear days (Pearson’s correlation, *r*=0.64, *p*<0.001 and *r*=0.59, *p*<0.001, respectively).

We found that the diversity of atmospheric microbiome on dusty days was significantly lower than on clear days (Mann–Whitney *U* test, *p*=0.007 and *p*=0.005, Figure 1C and 1F). This indicates that microbial communities in dusty atmospheric conditions were dominated by specific species and functions, resulting in a less even distribution of these features compared to clear atmospheric conditions. Furthermore, we did not observe a significant correlation between increasing PM_10_ concentrations and diversity specifically under dusty conditions (Pearson’s correlation, *r*=0.12, *p*=0.683 and *r*=0.25, *p*=0.410). This suggests that the dominance of specific species and functions in the microbiome during dusty conditions was not associated with the uplift of dust but rather influenced by potential ecological origins such as desert versus marine and local influences. In contrast, we observed that the diversity of atmospheric microbiome increased with rising temperatures throughout the sampling campaign for both taxonomic and functional attributes (Pearson’s correlation, *r*=0.35, *p*=0.019 and *r*=0.26 *p*=0.089, respectively). This trend was particularly pronounced on clear days (*r*=0.58, *p*<0.001 and *r*=0.32, *p*=0.069, respectively), although the increase in functional attributes did not reach statistical significance. These results imply that rising temperatures not only enhanced the observed number of species and functions but also contributed to a more uniform increase in their abundance.

In summary, environmental variables, particularly dust storms, and to a lesser extent, temperature fluctuations, significantly shape both the taxonomic and functional characteristics of the atmospheric microbiome, while air mass origin specifically influences taxonomic composition.

### Potential pathogens in the atmospheric microbiome

We next examined the presence of microbial species with pathogenic potential for human, animal, and plant health in air samples. Using metagenomic analysis, we identified DNA fragments matching a total of 117 potential pathogenic species. Bacteria constituted the majority, with 86 species, followed by fungi (25 species), other eukaryotes (4 species), and viruses (2 species). The bacterial species *Stenotrophomonas maltophilia* (3.15±0.27; mean±SD, N=45; prevalence), *Pasteurella multocida* (3.03±0.18, N=45) and *Klebsiella pneumonia* (2.89±0.30, N=45), and fungal species *Botrytis cinerea* (2.56±0.48, N=45), exhibited notably higher abundance (log_10_(CPM)) and prevalence across all samples compared to other species (Figure 2A). Details on the abundance of bacterial, fungal, eukaryotic, and viral species in each sample are provided in Supporting Table 5. Additional analysis methods and the rationale for taxonomic annotation using Kraken2 are available in the Supporting Information.

**Figure 2.**
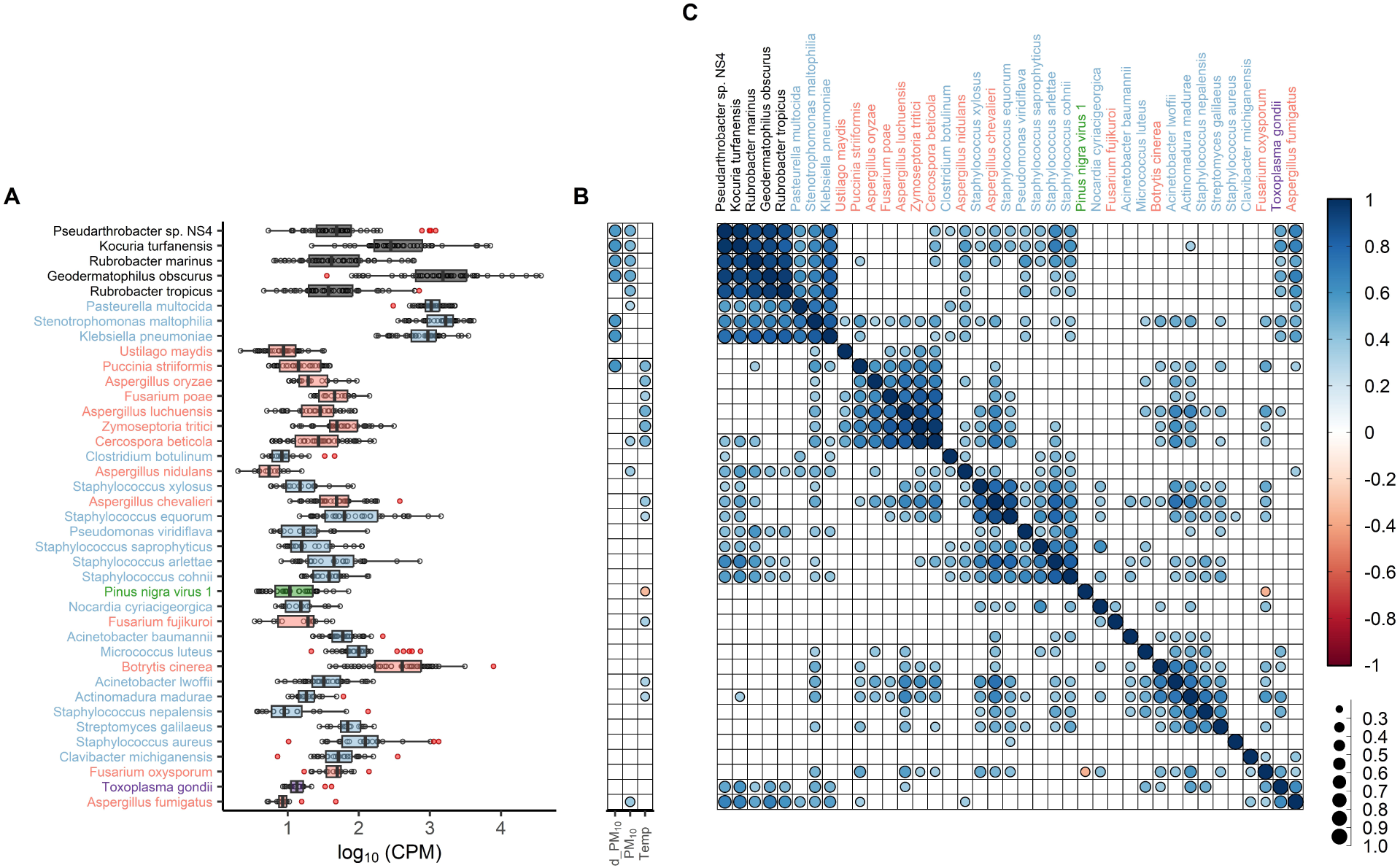
Relationships among potential pathogens, dust-indicator taxa, and environmental variables. (A) Abundance distribution (log_10_(CPM)) for dust-indicator taxa and prevalent potential pathogens. Boxplot details: center lines represent medians, hinges display the 25^th^ and 75^th^ percentiles, while whiskers extend to values within 1.5 times the Interquartile Range (IQR) from the hinges. Red-colored dots indicate outliers. (B) Pearson’s correlation analysis between individual taxa and environmental variables. ‘d_PM_10_’ signifies the correlation between daily mean PM_10_ concentration specifically within dusty atmospheric conditions, while ‘PM_10_’ denotes the correlation across all samples. ‘Temp’ represents the correlation between each taxon and the daily mean temperature. (C) Correlation matrix of Pearson’s coefficients showing co-occurrence patterns between potential pathogens and dust-indicator taxa. Correlations with an absolute Pearson’s coefficient (*r*) value above 0.3 and a P-value below 0.05, adjusted by the Benjamini–Hochberg method, were considered statistically significant. The visual representation uses color-coded correlation coefficient values and different-sized bins for ease of interpretation. Taxa names and boxplots are color-coded: black for dust-indicator taxa, light blue for bacteria, light red for fungi, green for viruses, and magenta for other eukaryotes. The x-axis and y-axis ordering were determined through hierarchical clustering using Ward’s method with the ward.D option in the ‘hclust’ package. This method minimizes within-cluster variance and combines clusters based on the smallest between-cluster distance.

Overall, we observed an increase in pathogen richness on dusty days, though this trend did not reach statistical significance (Pearson’s correlation, *r*=0.45, *p*=0.12), while pathogen diversity did not show a similar trend (*r*=0.22, *p*=0.46). However, both richness and diversity increased significantly with raising temperatures (*r*=0.50, *p*<0.001 and *r*=0.48, *p*<0.001, respectively), particularly on clear days (*r*=0.70, *p*<0.001 for both). This suggests that while dust storms may transport specific pathogens, higher temperatures enhance pathogen diversity, especially in clear conditions. These findings align with trends observed for all species in atmospheric microbial communities. Additionally, several bacterial pathogens such as *S. maltophilia*, *P. multocida* and *K. pneumonia* and fungal pathogens like *P. striiformis*, *Z. tritici* and *F. poae* showed correlation with increasing PM_10_ concentrations and temperatures (Figure 2B). However, the low abundance of these species and the inherent variability of the atmospheric microbiome made detecting strong associations with environmental variables challenging.

To gain further insight, we analyzed the co-occurrence patterns between taxa. We first identified dust-signature taxa that correlated strongly with rising PM_10_ concentrations: *G. obscurus* (Pearson’s correlation, *r*=0.51, *p*<0.001), *Pseudarthrobacter sp*. NS4 (*r*=0.48, *p*=0.001), *Rubrobacter marinus* (*r*=0.48, *p*=0.001), *Rubrobacter tropicus* (r=0.46, p=0.002), and *Kocuria turfanensis* (*r*=0.45, *p*=0.002). A literature review revealed that these taxa often inhabit harsh environments, such as desert rock varnish, and exhibit traits like UV-C resistance, further supporting their association with dusty conditions ^54–56^. We found that the dust-signature taxa showed strong correlations with each other, suggesting shared environmental origins. Moreover, while we identified only a few significant direct correlations between pathogenic taxa, PM_10_, and temperature, co-occurrence patterns indicated that several other pathogens were linked either with dust-signature taxa or with taxa correlated with higher temperatures (Figure 2C).

### Read mapping to the reference genomes

To validate these results, we mapped sequencing reads to the reference genomes of the 34 most prevalent potential pathogens and *G. obscurus*, which served as a reference due to its consistently high abundance throughout the sampling period. The primary objective of this analysis was to confirm whether the genomes of the identified pathogens were genuinely present or if the identifications were likely false positives. We achieved this by examining the genome coverage and read distribution. High genome coverage and a uniform distribution of reads across the reference genome would indicate the presence of these species in the air samples. Our analysis revealed varying levels of genome coverage (% of the genome’s base pairs covered by reads) and depth (the number of unique reads mapped to a specific nucleotide) for these species. For instance, reads from air samples covered approximately 30.0±19.8% (mean±SD) of the *G. obscurus* reference genome. In comparison, coverage for the most prevalent potential pathogens, despite their low abundance, was as follows: *S. maltophilia* (3.95±2.15%), *K. pneumonia* (2.38±3.16%), *B. cinerea* (1.54±2.29%), and *P. multocida* (0.84±0.22%). Additionally, the reads were uniformly distributed across the reference genomes, although with low mean depth (Figure 3 and Supporting Figure 13). Further details on genome coverage and depth for each species are available in Supporting Tables 6-8. These findings align with KrakenUniq results, indicating that Kraken’s classifications were based on marker k-mers distributed throughout the entire reference genome (Supporting Table 9 and Supporting Figure 14). Overall, the results confirm the accuracy of taxonomic classification and support the presence of these species in the air samples.

**Figure 3.**
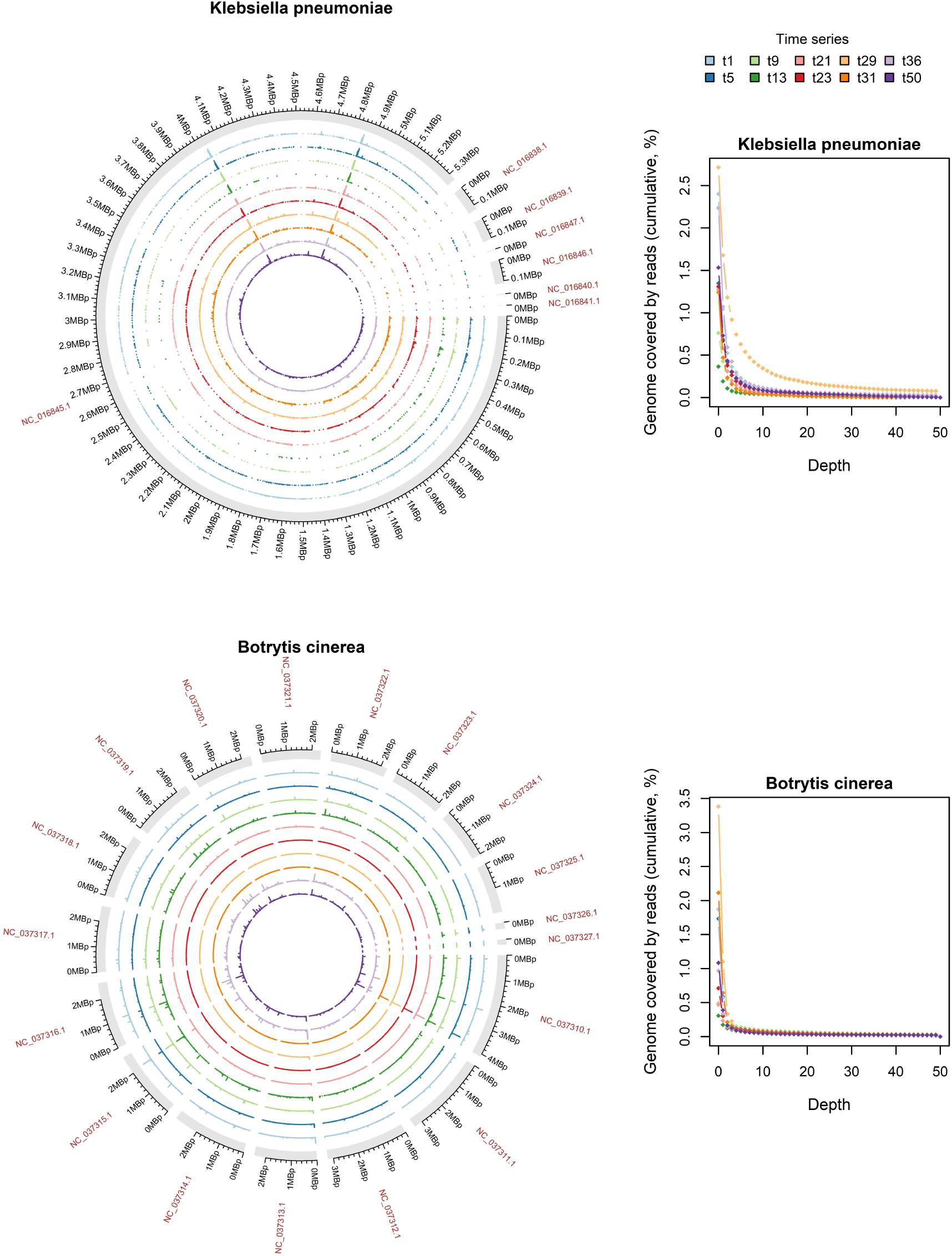
Read mapping results of individual samples to reference genomes, demonstrating coverage and depth. The circus plots show the read mapping results for the predominant bacterial pathogen *K. pneumoniae*, and the fungal pathogen *B. cinerea*. The outer circle represents the entire genome of each species, including chromosomes and plasmids, with a gap denoting the boundary. The RefSeq number is displayed at the top, outside the Circos plot, and genome size is annotated by numbers around each circle. Inside, colored circles (bar plots) depict air samples collected under various atmospheric conditions. For visual convenience, only five clear days (t5, t13, t23, t31 and t50) and five dusty days (t1, t9, t21, t29 and t36) are shown here. Each aligned read within a sample to the reference genome is depicted as a colored bar, indicating its specific position in the reference genome, with the height of the bar representing the depth. Coverage histograms illustrate the cumulative percentage of the total number of bases in the reference genome covered by different read depths.

### Antibiotic resistance and virulence-related genes

We identified genes that potentially confer resistance to 24 different classes of antibiotics, with varying abundance levels (Supporting Figure 15A and Supporting Table 10). Notably, genes associated with multidrug resistance, and resistance to bleomycin and fosfomycin, tetracycline, beta-lactam antibiotics resistance, and novobiocin and deoxycholate exhibited higher abundance (log_10_(RPKM)) and prevalence. Many of these genes also showed increased abundance as PM_10_ concentration and temperatures rose (Supporting Figure 15B). Similarly, we identified a wide range of virulence-related genes, each with varying abundance levels (Supporting Figure 16A and Supporting Table 11). These genes are linked to several protein domains, such as the RmlD substrate binding domain, virulence factor BrkB, lipid II flippase MurJ, and clp amino-terminal domain, all of which were more abundant and prevalent. Like the antibiotic resistance genes, their abundance increased with higher PM_10_ concentrations and elevated temperatures (Supporting Figure 16B). These protein domains are associated with virulence factors involved in cell adhesion, motility, biofilm formation, immunity, gene regulation, stress survival, and secretion systems. Some genes, such as the virulence factor BrkB, pneumococcal surface protein A (PspA), and the aerotolerance regulator N-terminal, are linked to dust storms, while others, like the PilZ domain and Clp amino-terminal domain, are associated with temperature variations. All these genes contribute to the virulence of respiratory pathogens ^57–68^. We also found that the overall richness of both antibiotic resistance and virulence-related genes increased with rising PM_10_ levels and temperatures. Notably, antibiotic resistance genes were more diverse on dusty days, while virulence-related genes exhibited greater diversity on clear days (Supporting Figures 17 and 18).

Next, we explored the host microorganisms harboring antibiotic resistance and virulence-related genes by assigning taxonomy to the contigs containing these genes. We found that 46.33±5.10% and 52.31±6.99% (mean ± SD) of all antibiotic resistance and virulence-contigs, respectively, did not classify with any level of taxa (Supporting Figures 19 and 20). This suggests two possibilities: either undiscovered species with these genes are prevalent in the atmospheric microbiome, or the taxonomic classification algorithm failed to identify certain taxa linked to these contigs. Additionally, we noted that only a small fraction of the antibiotic resistance and virulence-related contigs were associated with pathogenic species (Supporting Figures 21 and 22). We then investigated whether this was due to a technical limitation in the read assembly step. Given the low biomass, it is plausible that the DNA fragments of these less abundant species could not find matching overlapping reads. However, we found several assembled reads that were taxonomically classified as pathogenic species (Supporting Figure 23). For example, in six air samples, we detected a few contigs carrying antibiotic resistance genes that were classified as *K. pneumoniae*, a prevalent potential pathogen. These contigs had low abundance (RPKM: 3.78±2.42, mean±SD), while the total contig abundance of *K. pneumoniae* was substantially higher (271.57±218.12). Despite this, we did not find any virulence genes linked to *K. pneumoniae*. This suggests that the observed association between antibiotic resistance and virulence-related genes, and variations in PM_10_ and temperature, was not due to an increase in pathogen abundance. Moreover, we found that under dusty atmospheric conditions, most antibiotic resistance and virulence-related contigs were associated with dust-indicator and terrestrial taxa (Figure 4). Dust storms also exhibited significantly higher total abundance (RPKM) for these traits compared to samples from clear days. For instance, during a typical southwesterly dust storm (t7) and an easterly dust storm (t28), the total contig abundance for antibiotic resistance traits increased approximately 10- and 125-fold, respectively, compared to a typical clear day (t13), as shown in Figure 4. We also observed a 40-fold increase in resistance traits abundance when comparing clear-day air samples collected during lower temperatures (t13) to those collected during higher temperatures (t57). However, it remains unclear whether this increase is directly related to rising temperatures and their impact on microbial community composition, or to the cumulative effects of multiple dust storms occurring before the temperature rise. During this time of rising temperatures, we also recorded several daily temperature anomalies, likely due to heatwaves from neighboring regions, although these events were not associated with significant particle transport. It is also noteworthy that the southwesterly dust storm (t7) was sampled at the onset of the rainy, non-dusty season, with a major dust storm having occurred 3–4 months earlier. Intense precipitation during this period likely contributed to the wet removal of particulate matter and bioaerosols from the atmosphere.

**Figure 4.**
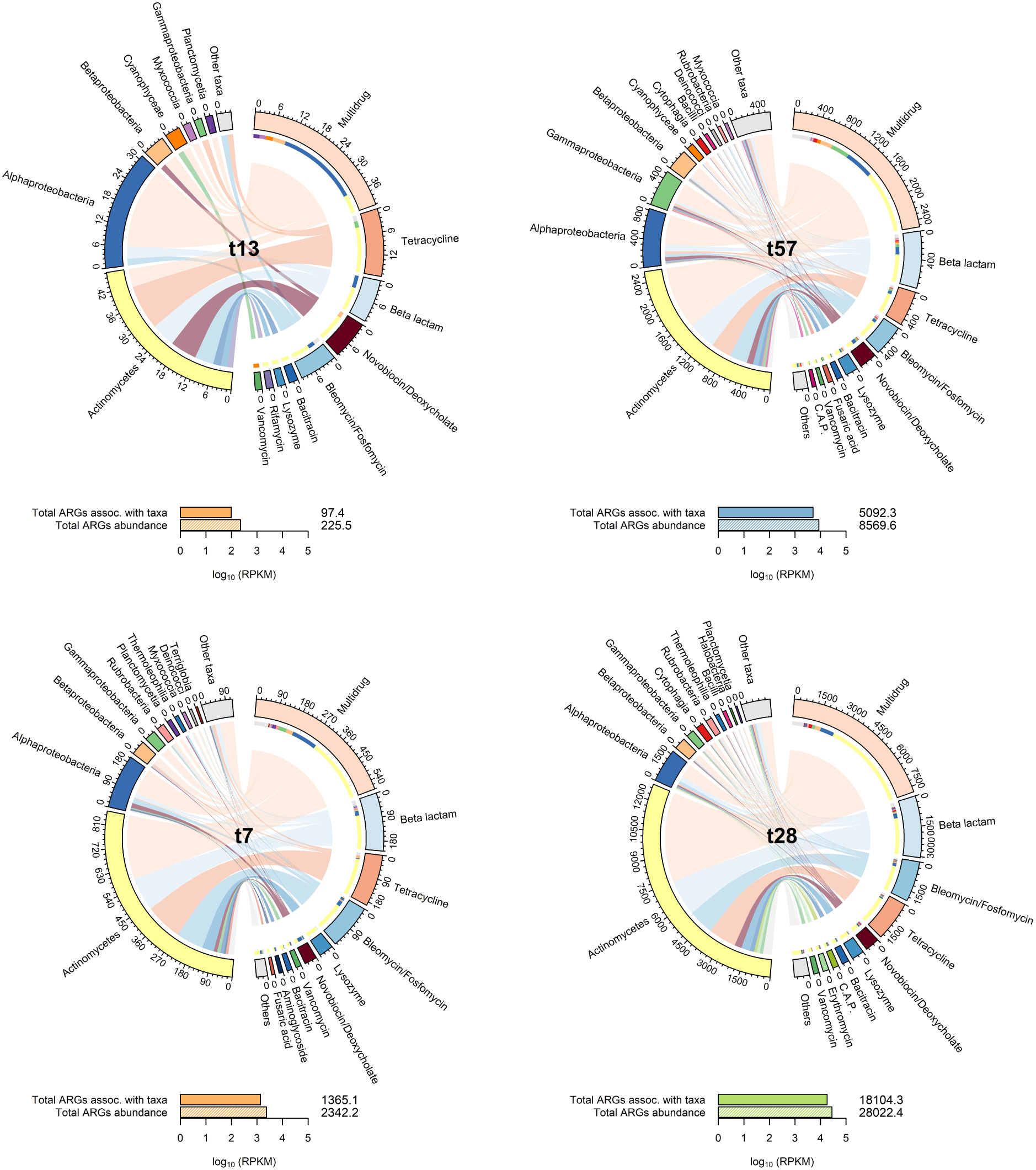
Host microorganisms carrying antibiotic resistance genes in air samples under clear and dusty atmospheric conditions. The RPKM values of contigs linked to both taxonomy and antibiotic resistance are labeled outside the circle. The sample name is displayed at the center of each chord diagram. For clarity, only four samples are shown: one from a clear day during a period of lower temperatures (t13), one from a clear day during a period of higher temperatures (t57), and two from dusty days corresponding to southwesterly (t7) and easterly (t28) dust storms. Inside the diagrams, colored sectors represent the top 10 taxa at the class level and the top 10 resistance types with the highest RPKM values. Gray sectors represent the sum of other taxa and functional features. Bar plots at the bottom of each chord diagram display the total log_10_(RPKM) values for all genes annotated with taxonomy and the total abundance of genes, including those linked to classified and unclassified taxa. The exact RPKM values are displayed above each bar. In the antibiotic resistance section, "C.A.P." stands for Cationic Antimicrobial Peptides.

## Discussion

The extensive metagenomic examination of the atmospheric microbiome in the Eastern Mediterranean region has provided novel insights into microbial dynamics and potential ecological impacts. Our findings reveal a diverse range of potential pathogens relevant to humans, animals, and plants, particularly associated with events of dust transport. For instance, certain bacterial species, such as *K. pneumonia* ^69^, *S. maltophilia* ^70^, and *P. multocida* ^71^ are specifically linked to human respiratory infections such as pneumonia. Fungal species like *A. fumigatus* ^72^, *A. nidulans* (which primarily affects immunocompromised individuals through conidia inhalation) ^73^ and *Aspergillus chevalieri* (with the primary potential to cause skin infections, notably cutaneous aspergillosis) ^74^ are associated with invasive pulmonary aspergillosis. Additionally, several *Staphylococcus* bacteria, including *Staphylococcus xylosus*, *Staphylococcus equorum*, *Staphylococcus arlettae*, *Staphylococcus cohnii*, and *Staphylococcus saprophyticus*, known to be associated with a range of infections such as bloodstream infections, urinary tract infections, and endocarditis ^75–79^. Moreover, the multi-host plant pathogen *Pseudomonas viridiflava*, with significant implications for agricultural crops such as tomatoes, melons, lettuce, chrysanthemums, and eggplants ^80^, is linked to dust storms. We also observed notable associations between temperature variations and certain potential pathogens, particularly fungi. For instance, *Zymoseptoria tritici*, *Fusarium poae*, and *P. striiformis*, can cause destructive wheat diseases like septoria tritici blotch, fusarium head blight and wheat yellow rust, respectively ^81–83^. Similarly, *C. beticola*, is the causative agent of the most detrimental foliar disease in sugar beet ^84^. In addition, certain potential pathogens associated with human diseases, such as *Aspergillus luchuensis*, known for causing invasive pulmonary aspergillosis ^85^, and *Aspergillus oryzae*, reported to induce peritonitis ^86^, exhibit a positive correlation with rising temperatures. While read mapping results further assure the presence of these species, we acknowledge the limitations of using shotgun metagenomic data to identify pathogens, particularly in low biomass, complex and diverse environmental microbiomes like those in air samples. This limitation is due to a reference database that is strongly biased towards human-associated organisms. Therefore, additional studies and methods can further validate the presence of specific pathogens identified in these air samples. These could include culture-based methods, although their efficacy may be compromised by the low biomass, and targeted qPCR, depending on the availability of species-specific primers.

The observed increasing trends in pathogen richness and compositional changes in abundance suggest a strong association between these species and environmental factors like dust storms and temperature fluctuations. This implies that dust storms could drive the dispersal of these microorganisms, while certain groups within the atmospheric microbiome appear to be temperature dependent. As previously discussed, many of these microorganisms are linked to human, animal, and plant diseases, raising concerns that dust storms and rising temperatures could pose threats to public and environmental health. However, further research is required to draw definitive conclusions about their immediate health implications.

These microorganisms inhabit diverse primary environments, including soil, oceans, humans, and other animals. Many are opportunistic pathogens, meaning they typically do not harm their hosts but can cause infections when the host’s immune system is compromised. Additionally, these pathogens are associated with a variety of diseases beyond those mentioned here. For instance, *P. multocida* has been reported to cause respiratory and invasive infections, particularly in immunocompromised individuals ^71,87,88^, but is more commonly associated with soft tissue infections following bites or scratches from domestic animals ^89^. Furthermore, the pathogenicity of these facultative pathogens is complex. It depends on various factors such as infective dosage, the host’s immune status, transmission route, and the presence of other competitive species on the host ^90,91^. For example, among these variables, the most critical factor is the initial dose of pathogens needed to trigger an infection in individual hosts, referred to as the infective dosage. Although data on inhalation-based infective doses for pathogens associated with dust storms and temperature variation is limited, previous research suggests that pathogens like *K. pneumoniae* and *S. maltophilia* generally have higher infective doses compared to others, such as *Shigella* ^92–95^. Therefore, even if the absolute abundance of these species increases under dusty conditions, as suggested by their rising relative abundance and overall higher microbial biomass, this does not necessarily indicate an immediate health or ecological risk. Moreover, although a prior study suggested that airborne bacteria might exhibit biochemical activity in dust samples ^15^, we lack direct evidence of the pathogen viability within the context of this study.

While we observed significant correlation between dust storms, rising temperatures, and specific antibiotic resistance and virulence genes, we identified only a small fraction of these genes on pathogen-associated contigs. This limitation likely arises from the technical challenges posed by low-biomass samples, which prevent the assembly of long enough contigs to reliably link these genes to specific pathogens. Co-assembling similar samples from the same origin could yield different results in this regard, warranting further investigation in future studies. Although the presence of antibiotic resistance and virulence genes in pathogens typically indicates an increased risk of pathogenicity, it is important to note that even when these genes are identified on pathogen contigs, they are insufficient on their own to cause disease. Other factors, such as gene expression, environmental conditions, and host susceptibility, are also necessary for an infection to occur. Thus, assessing the exact implications of dust storms and rising temperatures in this context remains complex.

We found that most of these genes, particularly on dusty days, were harbored by soil bacteria. This finding is not surprising, considering that soil microbiota represents one of the ancient evolutionary origins of antibiotic resistance and has been proposed as a reservoir of resistance genes ^96^. Although pristine soil microbiomes, such as those found in deserts, contain fewer antibiotic resistance genes compared to non-desert soils ^97^, it is plausible that we observed an increased abundance of these features on dusty days compared to clear days, which lack strong soil influence. Regarding virulence genes, it is reasonable to assume that microorganisms typically employ similar strategies to interact with hosts in microbial ecosystems. The emergence of new pathogens is often associated with gene loss or horizontal gene transfer from other species ^98,99^. For instance, pathogenic strains of *Enterococcus faecium* evolved from commensal species via horizontal gene transfer ^100^. Therefore, it is unsurprising that most virulence factors were connected to non-pathogenic microorganisms in this study. Additionally, some factors, such as those related to colonization, may assist these organisms in thriving in their habitat, thus possessing more generalized functions essential for survival in the environment ^101^.

We found that intense dust storms in the Middle East can carry antibiotic resistance genes, measured in RPKM abundance, at levels up to 125 times higher than those observed on clear days. The abundance of these genes on clear days in our study is consistent with previous findings from PM_10_ samples collected during clear days and those with several historically severe smog events in other locations ^35,102^. In contrast, the abundance of these genes on a typical dusty day in our study was notably higher than in environments like the human gut, ocean, anaerobic sludge, agricultural soil, cattle stool, drinking water and indoor air ^35,103–105^, where RPKM levels were similar to or lower than those in clear day air samples. Although differences in characterization methods between these studies and ours may introduce some bias, the consistent results from clear day air samples across studies enhance the reliability of these comparisons. Additionally, we observed that easterly dust storms typically exhibit higher levels of antibiotic resistance genes. This difference is likely due to the characteristics of the habitats from which these storms originate. Easterly storms traverse more anthropogenic regions, such as the Fertile Crescent, which spans northern Israel, Syria, and Iraq. These areas, heavily engaged in agriculture and vegetation, likely contribute to the elevated antibiotic resistance genes abundance in easterly air masses compared to those from the southwesterly and northwesterly directions. Conversely, southwesterly, and northwesterly air masses travel over the Mediterranean Sea, or the Sahara Desert, regions dominated by vast sand deserts and dry, sparsely populated areas. This observation aligns with a previous study on the global distribution of antibiotic resistance genes, which found that high-intensity human activity significantly increases the abundance of antibiotic resistance genes ^106^. Although the abundance of resistance genes in easterly dust storms was notably higher than in agricultural soil samples from another study ^35^, dust storms can accumulate and transport bioaerosols from a wide range of sources as they travel. Understanding the complexities of these interactions requires continuous air monitoring rather than relying on a brief two-month sampling period. Future studies should prioritize longer sampling campaigns that capture a broader range of conditions and sources.

While the lack of strong evidence connecting dust-borne pathogens to antibiotic resistance and virulence genes in our study suggests that the immediate health risks posed by these pathogens may be low, the dispersal of antibiotic resistance genes through dust storms could have indirect and long-term ecological consequences. First, our results indicate that dust storms likely come into direct contact with various emission sources, acquiring diverse functional traits, including resistance genes, as they travel through anthropogenic emissions. Second, the substantial genetic reservoir present in soil indicates a continuous emergence of new antibiotic resistance traits in this environment. Once these genes become mobile within microbial chromosomes, they have the potential to be transferred to pathogens. This transfer may occur within environmental settings or within the microbiota of humans or domestic animals. Even if these genes reside in inactive microbes or as extracellular DNA in air samples, they could still be assimilated into the genomes of viable cells, albeit with relatively low probabilities ^107^. Consequently, the dispersal of these genes through dust storms could facilitate the emergence of new resistance in pathogens across various ecosystems. Therefore, it is crucial to understand whether these genes are already mobile within the atmospheric microbiome, emphasizing the need for further studies in this area.

## Author Contributions

B.A.E. and Y.R. conceptualized and designed the study. B.A.E. collected air samples, extracted the DNA, and prepared them for sequencing, conducted bioinformatics and data analysis, interpreted, and visualized the results, and drafted the manuscript. B.A.E. and D.Z. jointly crafted the bioinformatics study design. Y.R. and D.Z. contributed to reviewing and editing the manuscript. All authors actively participated in result discussions and approved the final manuscript.

## References

1. Fröhlich-Nowoisky, J. et al. Bioaerosols in the Earth system: Climate, health, and ecosystem interactions. Atmospheric Research 182, 346–376 (2016). 10.1016/j.atmosres.2016.07.018

2. Failor, K. C., Schmale, D. G., Vinatzer, B. A. & Monteil, C. L. Ice nucleation active bacteria in precipitation are genetically diverse and nucleate ice by employing different mechanisms. The ISME Journal 11, 2740–2753 (2017). 10.1038/ismej.2017.124

3. de Araujo, G. G., Rodrigues, F., Gonçalves, F. L. T. & Galante, D. Survival and ice nucleation activity of Pseudomonas syringae strains exposed to simulated high-altitude atmospheric conditions. Scientific Reports 9, 7768 (2019). 10.1038/s41598-019-44283-3

4. Bauer, H. et al. Airborne bacteria as cloud condensation nuclei. Journal of Geophysical Research: Atmospheres 108 (2003). 10.1029/2003JD003545

5. Amato, P. et al. Metatranscriptomic exploration of microbial functioning in clouds. Scientific Reports 9, 4383 (2019). 10.1038/s41598-019-41032-4

6. Vaïtilingom, M. et al. Potential impact of microbial activity on the oxidant capacity and organic carbon budget in clouds. Proceedings of the National Academy of Sciences 110, 559–564 (2013). doi:10.1073/pnas.1205743110

7. Kobziar, L. N. et al. Wildland fire smoke alters the composition, diversity, and potential atmospheric function of microbial life in the aerobiome. ISME Commun 2, 8 (2022). 10.1038/s43705-022-00089-5

8. Joung, Y. S., Ge, Z. & Buie, C. R. Bioaerosol generation by raindrops on soil. Nat Commun 8, 14668 (2017). 10.1038/ncomms14668

9. Michaud, J. M. et al. Taxon-specific aerosolization of bacteria and viruses in an experimental ocean-atmosphere mesocosm. Nat Commun 9, 2017 (2018). 10.1038/s41467-018-04409-z

10. Varga, G., Újvári, G. & Kovács, J. Spatiotemporal patterns of Saharan dust outbreaks in the Mediterranean Basin. Aeolian Research 15, 151–160 (2014). 10.1016/j.aeolia.2014.06.005

11. Bodenheimer, S., Lensky, I. M. & Dayan, U. Characterization of Eastern Mediterranean dust storms by area of origin; North Africa vs. Arabian Peninsula. Atmospheric Environment 198, 158–165 (2019). 10.1016/j.atmosenv.2018.10.034

12. Mazar, Y., Cytryn, E., Erel, Y. & Rudich, Y. Effect of Dust Storms on the Atmospheric Microbiome in the Eastern Mediterranean. Environ Sci Technol 50, 4194–4202 (2016). 10.1021/acs.est.5b06348

13. Gat, D., Mazar, Y., Cytryn, E. & Rudich, Y. Origin-Dependent Variations in the Atmospheric Microbiome Community in Eastern Mediterranean Dust Storms. Environ Sci Technol 51, 6709–6718 (2017). 10.1021/acs.est.7b00362

14. Mayol, E. et al. Long-range transport of airborne microbes over the global tropical and subtropical ocean. Nat Commun 8, 201 (2017). 10.1038/s41467-017-00110-9

15. Erkorkmaz, B. A., Gat, D. & Rudich, Y. Aerial transport of bacteria by dust plumes in the Eastern Mediterranean revealed by complementary rRNA/rRNA-gene sequencing. Communications Earth & Environment 4, 24 (2023). 10.1038/s43247-023-00679-8

16. Rahav, E., Ovadia, G., Paytan, A. & Herut, B. Contribution of airborne microbes to bacterial production and N2 fixation in seawater upon aerosol deposition. Geophysical Research Letters 43, 719–727 (2016). 10.1002/2015gl066898

17. Guo, C. et al. Shifts in Microbial Community Structure and Activity in the Ultra-Oligotrophic Eastern Mediterranean Sea Driven by the Deposition of Saharan Dust and European Aerosols. Frontiers in Marine Science 3 (2016). 10.3389/fmars.2016.00170

18. Lang-Yona, N. et al. Annual distribution of allergenic fungal spores in atmospheric particulate matter in the Eastern Mediterranean; a comparative study between ergosterol and quantitative PCR analysis. Atmospheric Chemistry and Physics 12, 2681–2690 (2012). 10.5194/acp-12-2681-2012

19. Brodsky, H. et al. Assessing long-distance atmospheric transport of soilborne plant pathogens. Environ Res Lett 18 (2023). ARTN 104021 10.1088/1748-9326/acf50c

20. Gonzalez-Martin, C., Teigell-Perez, N., Valladares, B. & Griffin, D. W. The Global Dispersion of Pathogenic Microorganisms by Dust Storms and Its Relevance to Agriculture. Adv Agron 127, 1–41 (2014). 10.1016/B978-0-12-800131-8.00001-7

21. Rotem, J. Sand and dust storms as factors leading to alternaria blight epidemics on potatoes and tomatoes. Agricultural Meteorology 2, 281–288 (1965). 10.1016/0002-1571(65)90014-2

22. Zhang, C. et al. Mortality risks from a spectrum of causes associated with sand and dust storms in China. Nature Communications 14, 6867 (2023). 10.1038/s41467-023-42530-w

23. Sajani, S. Z. et al. Saharan dust and daily mortality in Emilia-Romagna (Italy). Occupational and Environmental Medicine 68, 446–451 (2011). 10.1136/oem.2010.058156

24. Meng, Z. & Lu, B. Dust events as a risk factor for daily hospitalization for respiratory and cardiovascular diseases in Minqin, China. Atmospheric Environment 41, 7048–7058 (2007). 10.1016/j.atmosenv.2007.05.006

25. Chan, C.-C. & Ng, H.-C. A case-crossover analysis of Asian dust storms and mortality in the downwind areas using 14-year data in Taipei. Science of The Total Environment 410-411, 47–52 (2011). 10.1016/j.scitotenv.2011.09.031

26. Tam, W. W. S., Wong, T. W., Wong, A. H. S. & Hui, D. S. C. Effect of dust storm events on daily emergency admissions for respiratory diseases. Respirology 17, 143–148 (2012). 10.1111/j.1440-1843.2011.02056.x

27. Vodonos, A. et al. The impact of desert dust exposures on hospitalizations due to exacerbation of chronic obstructive pulmonary disease. Air Quality, Atmosphere & Health 7, 433–439 (2014). 10.1007/s11869-014-0253-z

28. Lorentzou, C. et al. Extreme desert dust storms and COPD morbidity on the island of Crete. Int J Chron Obstruct Pulmon Dis 14, 1763–1768 (2019). 10.2147/copd.S208108

29. Zittis, G. et al. Climate Change and Weather Extremes in the Eastern Mediterranean and Middle East. Reviews of Geophysics 60, e2021RG000762 (2022). 10.1029/2021rg000762

30. Ozturk, T., Turp, M. T., Türkeş, M. & Kurnaz, M. L. Future projections of temperature and precipitation climatology for CORDEX-MENA domain using RegCM4.4. Atmospheric Research 206, 87–107 (2018). 10.1016/j.atmosres.2018.02.009

31. Luhung, I. et al. Experimental parameters defining ultra-low biomass bioaerosol analysis. NPJ Biofilms Microbiomes 7, 37 (2021). 10.1038/s41522-021-00209-4

32. Gat, D. et al. Size-Resolved Community Structure of Bacteria and Fungi Transported by Dust in the Middle East. Front Microbiol 12, 744117 (2021). 10.3389/fmicb.2021.744117

33. Gusareva, E. S. et al. Microbial communities in the tropical air ecosystem follow a precise diel cycle. Proc Natl Acad Sci U S A 116, 23299–23308 (2019). 10.1073/pnas.1908493116

34. Amato, P. et al. Metatranscriptomic exploration of microbial functioning in clouds. Sci Rep 9, 4383 (2019). 10.1038/s41598-019-41032-4

35. Qin, N. et al. Longitudinal survey of microbiome associated with particulate matter in a megacity. Genome Biology 21, 55 (2020). 10.1186/s13059-020-01964-x

36. Gat, D., Zimmermann, R. & Rudich, Y. Functional Genes Profile of Atmospheric Dust in the East Mediterranean Suggests Widespread Anthropogenic Influence on Aerobiome Composition. Journal of Geophysical Research: Biogeosciences 127, e2022JG007022 (2022). 10.1029/2022JG007022

37. Peng, X., Gat, D., Paytan, A. & Rudich, Y. The Response of Airborne Mycobiome to Dust Storms in the Eastern Mediterranean. Journal of Fungi 7, 802 (2021).

38. Bolger, A. M., Lohse, M. & Usadel, B. Trimmomatic: a flexible trimmer for Illumina sequence data. Bioinformatics 30, 2114–2120 (2014). 10.1093/bioinformatics/btu170

39. Wood, D. E., Lu, J. & Langmead, B. Improved metagenomic analysis with Kraken 2. Genome Biol 20, 257 (2019). 10.1186/s13059-019-1891-0

40. Lu, J., Breitwieser, F. P., Thielen, P. & Salzberg, S. L. Bracken: estimating species abundance in metagenomics data. Peerj Comput Sci 3, e104 (2017). 10.7717/peerj-cs.104

41. Breitwieser, F. P., Baker, D. N. & Salzberg, S. L. KrakenUniq: confident and fast metagenomics classification using unique k-mer counts. Genome Biol 19, 198 (2018). 10.1186/s13059-018-1568-0

42. Langmead, B. & Salzberg, S. L. Fast gapped-read alignment with Bowtie 2. Nature Methods 9, 357–359 (2012). 10.1038/nmeth.1923

43. Danecek, P. et al. Twelve years of SAMtools and BCFtools. GigaScience 10 (2021). 10.1093/gigascience/giab008

44. Quinlan, A. R. & Hall, I. M. BEDTools: a flexible suite of utilities for comparing genomic features. Bioinformatics 26, 841–842 (2010). 10.1093/bioinformatics/btq033

45. Gu, Z., Gu, L., Eils, R., Schlesner, M. & Brors, B. circlize implements and enhances circular visualization in R. Bioinformatics 30, 2811–2812 (2014). 10.1093/bioinformatics/btu393

46. Buchfink, B., Xie, C. & Huson, D. H. Fast and sensitive protein alignment using DIAMOND. Nat Methods 12, 59–60 (2015). 10.1038/nmeth.3176

47. Huson, D. H., Auch, A. F., Qi, J. & Schuster, S. C. MEGAN analysis of metagenomic data. Genome Res 17, 377–386 (2007). 10.1101/gr.5969107

48. Nurk, S., Meleshko, D., Korobeynikov, A. & Pevzner, P. A. metaSPAdes: a new versatile metagenomic assembler. Genome Res 27, 824–834 (2017). 10.1101/gr.213959.116

49. Hyatt, D. et al. Prodigal: prokaryotic gene recognition and translation initiation site identification. BMC Bioinformatics 11, 119 (2010). 10.1186/1471-2105-11-119

50. Eddy, S. R. A new generation of homology search tools based on probabilistic inference. Genome Inform 23, 205–211 (2009).

51. Mistry, J. et al. Pfam: The protein families database in 2021. Nucleic Acids Res 49, D412–D419 (2021). 10.1093/nar/gkaa913

52. Shen, W. & Ren, H. TaxonKit: A practical and efficient NCBI taxonomy toolkit. Journal of Genetics and Genomics 48, 844–850 (2021). 10.1016/j.jgg.2021.03.006

53. Battistuzzi, F. U. & Hedges, S. B. A major clade of prokaryotes with ancient adaptations to life on land. Mol Biol Evol 26, 335–343 (2009). 10.1093/molbev/msn247

54. Ivanova, N. et al. Complete genome sequence of Geodermatophilus obscurus type strain (G-20). Stand Genomic Sci 2, 158–167 (2010). 10.4056/sigs.711311

55. Chen, R. W. et al. Rubrobacter tropicus sp. nov. and Rubrobacter marinus sp. nov., isolated from deep-sea sediment of the South China Sea. Int J Syst Evol Microbiol 70, 5576–5585 (2020). 10.1099/ijsem.0.004449

56. Zhou, G. et al. Kocuria flava sp. nov. and Kocuria turfanensis sp. nov., airborne actinobacteria isolated from Xinjiang, China. Int J Syst Evol Microbiol 58, 1304–1307 (2008). 10.1099/ijs.0.65323-0

57. Fernandez, R. C. & Weiss, A. A. Cloning and sequencing of a Bordetella pertussis serum resistance locus. Infect Immun 62, 4727–4738 (1994). 10.1128/iai.62.11.4727-4738.1994

58. McCool, T. L., Cate, T. R., Moy, G. & Weiser, J. N. The immune response to pneumococcal proteins during experimental human carriage. J Exp Med 195, 359–365 (2002). 10.1084/jem.20011576

59. Mukerji, R. et al. Pneumococcal surface protein A inhibits complement deposition on the pneumococcal surface by competing with the binding of C-reactive protein to cell-surface phosphocholine. J Immunol 189, 5327–5335 (2012). 10.4049/jimmunol.1201967

60. Park, S. S. et al. Streptococcus pneumoniae Binds to Host Lactate Dehydrogenase via PspA and PspC To Enhance Virulence. mBio 12, 10.1128/mbio.00673-00621 (2021). https://doi.org:10.1128/mBio.00673-21

61. Lafontaine, E. R. et al. The autotransporter protein BatA is a protective antigen against lethal aerosol infection with Burkholderia mallei and Burkholderia pseudomallei. Vaccine X 1, 100002 (2019). 10.1016/j.jvacx.2018.100002

62. Yesilkaya, H., Andisi, V. F., Andrew, P. W. & Bijlsma, J. J. Streptococcus pneumoniae and reactive oxygen species: an unusual approach to living with radicals. Trends Microbiol 21, 187–195 (2013). 10.1016/j.tim.2013.01.004

63. Schumacher, M. A. & Zeng, W. Structures of the activator of K. pneumonia biofilm formation, MrkH, indicates PilZ domains involved in c-di-GMP and DNA binding. Proc Natl Acad Sci U S A 113, 10067–10072 (2016). 10.1073/pnas.1607503113

64. Wilksch, J. J. et al. MrkH, a novel c-di-GMP-dependent transcriptional activator, controls Klebsiella pneumoniae biofilm formation by regulating type 3 fimbriae expression. PLoS Pathog 7, e1002204 (2011). 10.1371/journal.ppat.1002204

65. Porankiewicz, J., Wang, J. & Clarke, A. K. New insights into the ATP-dependent Clp protease: Escherichia coli and beyond. Mol Microbiol 32, 449–458 (1999). 10.1046/j.1365-2958.1999.01357.x

66. Kannan, T. R., Musatovova, O., Gowda, P. & Baseman, J. B. Characterization of a unique ClpB protein of Mycoplasma pneumoniae and its impact on growth. Infect Immun 76, 5082–5092 (2008). 10.1128/IAI.00698-08

67. Tripathi, P., Singh, L. K., Kumari, S., Hakiem, O. R. & Batra, J. K. ClpB is an essential stress regulator of Mycobacterium tuberculosis and endows survival advantage to dormant bacilli. Int J Med Microbiol 310, 151402 (2020). 10.1016/j.ijmm.2020.151402

68. Frees, D. et al. Clp ATPases are required for stress tolerance, intracellular replication and biofilm formation in Staphylococcus aureus. Mol Microbiol 54, 1445–1462 (2004). 10.1111/j.1365-2958.2004.04368.x

69. Holt, K. E. et al. Genomic analysis of diversity, population structure, virulence, and antimicrobial resistance in Klebsiella pneumoniae, an urgent threat to public health. Proc Natl Acad Sci U S A 112, E3574–3581 (2015). 10.1073/pnas.1501049112

70. Brooke, J. S. Stenotrophomonas maltophilia: an emerging global opportunistic pathogen. Clin Microbiol Rev 25, 2–41 (2012). 10.1128/CMR.00019-11

71. Pak, S., Valencia, D., Decker, J., Valencia, V. & Askaroglu, Y. Pasteurella multocida pneumonia in an immunocompetent patient: Case report and systematic review of literature. Lung India 35, 237–240 (2018). 10.4103/lungindia.lungindia_482_17

72. Kousha, M., Tadi, R. & Soubani, A. O. Pulmonary aspergillosis: a clinical review. Eur Respir Rev 20, 156–174 (2011). 10.1183/09059180.00001011

73. Bastos, R. W. et al. Functional Characterization of Clinical Isolates of the Opportunistic Fungal Pathogen Aspergillus nidulans. mSphere 5, 10.1128/msphere.00153-00120 (2020). https://doi.org:10.1128/mSphere.00153-20

74. Naidu, J. & Singh, S. M. Aspergillus chevalieri (Mangin) Thom and Church: a new opportunistic pathogen of human cutaneous aspergillosis. Mycoses 37, 271–274 (1994). 10.1111/j.1439-0507.1994.tb00425.x

75. Giordano, N. et al. Erythema nodosum associated with Staphylococcus xylosus septicemia. J Microbiol Immunol Infect 49, 134–137 (2016). 10.1016/j.jmii.2012.10.003

76. Novakova, D. et al. Staphylococcus equorum and Staphylococcus succinus isolated from human clinical specimens. J Med Microbiol 55, 523–528 (2006). 10.1099/jmm.0.46246-0

77. Lavecchia, A. et al. Staphylococcus arlettae Genomics: Novel Insights on Candidate Antibiotic Resistance and Virulence Genes in an Emerging Opportunistic Pathogen. Microorganisms 7 (2019). 10.3390/microorganisms7110580

78. Motta, J. C., Forero-Carreno, C., Arango, A. & Sanchez, M. Staphylococcus cohnii endocarditis in native valve. New Microbes New Infect 38, 100825 (2020). 10.1016/j.nmni.2020.100825

79. Hur, J. et al. Staphylococcus saprophyticus Bacteremia originating from Urinary Tract Infections: A Case Report and Literature Review. Infect Chemother 48, 136–139 (2016). 10.3947/ic.2016.48.2.136

80. Sarris, P. F., Trantas, E. A., Mpalantinaki, E., Ververidis, F. & Goumas, D. E. Pseudomonas viridiflava, a multi host plant pathogen with significant genetic variation at the molecular level. PLoS One 7, e36090 (2012). 10.1371/journal.pone.0036090

81. Fones, H. & Gurr, S. The impact of Septoria tritici Blotch disease on wheat: An EU perspective. Fungal Genet Biol 79, 3–7 (2015). 10.1016/j.fgb.2015.04.004

82. Xu, X. M. et al. Predominance and association of pathogenic fungi causing Fusarium ear blightin wheat in four European countries. European Journal of Plant Pathology 112, 143–154 (2005). 10.1007/s10658-005-2446-7

83. Chen, W., Wellings, C., Chen, X., Kang, Z. & Liu, T. Wheat stripe (yellow) rust caused by Puccinia striiformis f. sp. tritici. Mol Plant Pathol 15, 433–446 (2014). 10.1111/mpp.12116

84. Rangel, L. I. et al. Cercospora beticola: The intoxicating lifestyle of the leaf spot pathogen of sugar beet. Mol Plant Pathol 21, 1020–1041 (2020). 10.1111/mpp.12962

85. Wang, Q. et al. Triazole-resistant Aspergillus luchuensis, an industrially important black Aspergillus spp. used in fermentation in East Asia, isolated from the patient with invasive pulmonary aspergillosis in China. Emerg Microbes Infect 11, 1435–1438 (2022). 10.1080/22221751.2022.2076614

86. Schwetz, I. et al. Aspergillus oryzae peritonitis in CAPD: case report and review of the literature. Am J Kidney Dis 49, 701–704 (2007). 10.1053/j.ajkd.2007.02.260

87. Yadav, S. A Case of Pneumonia Caused by Pasteurella multocida in an Immunocompetent Indian Male. Cureus 14, e28820 (2022). 10.7759/cureus.28820

88. Kofteridis, D. P. et al. Bacteremic community-acquired pneumonia due to Pasteurella multocida. International Journal of Infectious Diseases 13, e81–e83 (2009). 10.1016/j.ijid.2008.06.023

89. Wilson, B. A. & Ho, M. Pasteurella multocida: from zoonosis to cellular microbiology. Clin Microbiol Rev 26, 631–655 (2013). 10.1128/cmr.00024-13

90. Keen, E. C. Paradigms of pathogenesis: targeting the mobile genetic elements of disease. Front Cell Infect Microbiol 2, 161 (2012). 10.3389/fcimb.2012.00161

91. Fierer, J., Looney, D. & Pechère, J.-C. in Infectious Diseases (Fourth Edition) (eds Jonathan Cohen, William G. Powderly, & Steven M. Opal) 4–25.e21 (Elsevier, 2017).

92. Malina, J., Hofmann, J. & Franĕk, J. Informative value of a mouse model of Klebsiella pneumoniae infection used as a host-resistance assay. Folia Microbiol (Praha) 36, 183–191 (1991). 10.1007/bf02814501

93. Lawlor, M. S., Hsu, J., Rick, P. D. & Miller, V. L. Identification of Klebsiella pneumoniae virulence determinants using an intranasal infection model. Molecular Microbiology 58, 1054–1073 (2005). 10.1111/j.1365-2958.2005.04918.x

94. Pompilio, A. et al. Stenotrophomonas maltophilia Phenotypic and Genotypic Diversity during a 10-year Colonization in the Lungs of a Cystic Fibrosis Patient. Frontiers in Microbiology 7 (2016). 10.3389/fmicb.2016.01551

95. DuPont, H. L., Levine, M. M., Hornick, R. B. & Formal, S. B. Inoculum size in shigellosis and implications for expected mode of transmission. J Infect Dis 159, 1126–1128 (1989). 10.1093/infdis/159.6.1126

96. Forsberg, K. J. et al. The Shared Antibiotic Resistome of Soil Bacteria and Human Pathogens. Science 337, 1107–1111 (2012). doi:10.1126/science.1220761

97. Fierer, N. et al. Cross-biome metagenomic analyses of soil microbial communities and their functional attributes. Proceedings of the National Academy of Sciences 109, 21390–21395 (2012). doi:10.1073/pnas.1215210110

98. Pallen, M. J. & Wren, B. W. Bacterial pathogenomics. Nature 449, 835–842 (2007). 10.1038/nature06248

99. Brown, N. F., Wickham, M. E., Coombes, B. K. & Finlay, B. B. Crossing the line: selection and evolution of virulence traits. PLoS Pathog 2, e42 (2006). 10.1371/journal.ppat.0020042

100. Leavis, H. L. et al. Insertion Sequence–Driven Diversification Creates a Globally Dispersed Emerging Multiresistant Subspecies of E. faecium. PLOS Pathogens 3, e7 (2007). 10.1371/journal.ppat.0030007

101. Niu, C. et al. Common and pathogen-specific virulence factors are different in function and structure. Virulence 4, 473–482 (2013). 10.4161/viru.25730

102. Pal, C., Bengtsson-Palme, J., Kristiansson, E. & Larsson, D. G. J. The structure and diversity of human, animal and environmental resistomes. Microbiome 4, 54 (2016). 10.1186/s40168-016-0199-5

103. Ben Maamar, S., et al. Mobilizable antibiotic resistance genes are present in dust microbial communities. PLOS Pathogens 16, e1008211 (2020). 10.1371/journal.ppat.1008211

104. Kang, K. et al. The Environmental Exposures and Inner- and Intercity Traffic Flows of the Metro System May Contribute to the Skin Microbiome and Resistome. Cell Reports 24, 1190–1202.e1195 (2018). 10.1016/j.celrep.2018.06.109

105. Hartmann, E. M. et al. Antimicrobial Chemicals Are Associated with Elevated Antibiotic Resistance Genes in the Indoor Dust Microbiome. Environmental Science & Technology 50, 9807–9815 (2016). 10.1021/acs.est.6b00262

106. Zhang, Z. et al. Assessment of global health risk of antibiotic resistance genes. Nat Commun 13, 1553 (2022). 10.1038/s41467-022-29283-8

107. Larsson, D. G. J. & Flach, C.-F. Antibiotic resistance in the environment. Nature Reviews Microbiology 20, 257–269 (2022). 10.1038/s41579-021-00649-x

