## Supporting Information for "Dust storm-driven dispersal of potential pathogens and antibiotic resistance genes in the Eastern Mediterranean"

Burak A. Erkorkmaz *et al*.

**This PDF file includes:**

Supplementary Text

Supporting Figures 1 to 24

Supporting Tables 1 to 13

Supporting Movies 1 to 4

**Other Supporting Information for this manuscript include the following:**

Supporting Tables 1 to 3 and 5 to 13 (separate file)

**Supplementary Text**

**Materials and Methods**

**Sample collection**

To predict dust storms, we utilized various atmospheric forecast platforms, including: Forecast Maps of Dust by the University of Athens (https://forecast.uoa.gr/), the AEMET Dust Forecast platform (https://dust.aemet.es/), Copernicus Atmosphere Monitoring Service (https://atmosphere.copernicus.eu/) and Windy (https://www.windy.com/). We verified the accuracy of dust storm predictions by cross-referencing them with $PM_{10}$ data obtained from the Rehovot Air Monitoring station of the Israeli Ministry of Environmental Protection (https://air.sviva.gov.il/). To mitigate potential confounding effect of the diel cycle of the atmospheric microbiome [1], we collected all samples specifically from 9:00 to 12:00. We chose this time frame to minimize variations in microbial composition and activity throughout the day. However, in cases where dust forecasts indicated a dust storm or other significant atmospheric events outside this time slot, we adjusted sample collection accordingly to ensure representative sampling.

For metagenomics analysis, we employed the Coriolis µ microbial air sampler (Bertin Technologies) to collect air samples for 2 hours, at a flow rate of 200 L/min. The Coriolis µ microbial air sampler captures particles larger than 0.5 µm, with a collection efficiency with a $D_{50}$ value, denoting the particle size at which the collection efficiency reaches 50%, exceeding 0.5 µm. During sampling, we collected particulate matter and bioaerosols in 15 ml of a nucleic acids preservation solution to maintain sample integrity [2]. To prevent evaporation, we supplemented the samples with molecular-grade water at a flow rate of 0.5-1.5 ml per minute. Before each sampling event, we thoroughly decontaminate the Coriolis sampler as detailly described below. Following collection, we concentrated the sample onto autoclaved and sterilized polyethersulfone (PES) filters (diameter of 25 mm and a pore size of 0.1 µm (Sartorius Stedim Biotech)) using a sterile single-use syringe and reusable syringe filter holders (Sartorius Stedim Biotech). Then, we transferred the filter samples into Falcon tubes and stored at -20°C for preservation. We conducted DNA extraction from the filters within 2-3 days after collection to ensure optimal sample quality. To generate operating blanks, we replicated the same procedure, but we operated the sampler for 5 seconds. Consequently, we obtained a total of seven operating blanks on randomly selected dates before the actual sample collection process. This ensured the capture of background environmental conditions and potential contamination present in the sampling setup.

**Decontamination of Coriolis sampler**

To ensure thorough cleaning and removal of potential contaminants, we conducted a series of meticulous steps for the decontamination of the Coriolis sampler before each sampling event. Initially, we soaked the cane, air intake, and evaporation tubing in a 1.5% sodium hypochlorite (NaClO) solution for 15 minutes. Following this, we rinsed them with ultrapure water, washed them with 70% ethanol (EtOH), and finally rinsed them again with ultrapure water to achieve complete decontamination. For the reusable sampling cones and filter holders of the Coriolis sampler, we adopted a specific decontamination procedure. We began by washing the cones and filter holders with tap water and then immersing them in a 3% NaClO solution for 15 minutes to ensure effective disinfection. After the NaClO soak, we thoroughly rinsed the cones and filter holders with ultrapure water to eliminate any residual disinfectant. Subsequently, we sterilized the surfaces further by washing them for 10-15 seconds with 96% EtOH. Finally, we thoroughly rinsed the cones and filter holders with ultrapure water and autoclaved them at 121°C for 15 minutes. To validate the efficacy of this decontamination procedure, we conducted qPCR analysis using previously described primers and methods [2]. The results confirmed the absence of 16S ribosomal DNA and RNA, thus verifying the effectiveness of the decontamination process.

**Collection of meteorological data and dust column density maps**

We collected data on particulate matter concentration (PM, $\mu g\ m^{-3}$), temperature (T, °C), relative humidity (RH, %), and rain (mm) from the Rehovot Air Monitoring station, located approximately 1 km from the sampling site. This station is part of the Israeli Ministry of Environmental Protection network. Meteorological data were recorded in 5-min time intervals. To estimate the origin of the dust, we acquired time-averaged maps of dust column mass density (at a $0.5° \times 0.625°$ resolution) and reanalysis meteorological data from the Modern-Era Retrospective analysis for Research and Applications (MERRA-2) for each sampling event. For each event, we performed a retrospective analysis using the Giovanni online data system, developed and maintained by the NASA GES DISC [3], by using hourly frames of dust column density maps 72 hours prior to the sampling event. These frames were compiled into dust column density map movies using the 'av' package in R, with data at 6-hour intervals. Additionally, we investigated the origin of the air mass with the Hybrid Single-Particle Lagrangian Integrated Trajectory model (HYSPLIT) (https://www.ready.noaa.gov/index.php/) [4,5]. For each analysis, we calculated a 72-hour back trajectory at three distinct altitudes (0, 50, and 100 meters above ground level).

**List of known pathogenic microorganisms**

To assess the presence of potential human, animal, and plant pathogens within the atmospheric microbiome, we compiled a list of known pathogenic microorganisms from various publicly available databases (details can be found in Supporting Table 12). For pathogen names provided at the genus level, we performed a comprehensive literature search to confirm any pathogenic connections for all species detected within the metagenomic samples corresponding to the listed genera in the pathogen database.

**Data processing and metagenomic analysis**

We performed taxonomic classification of the metagenomic data using Kraken2 (v2.1.2), which employs exact k-mer matching directly on paired reads (--paired --minimum-hit-groups 10 --confidence 0.1) [6]. In this procedure, we constructed a custom database of taxa-specific k-mers using complete reference genomes/proteins from diverse taxonomic groups, including archaea, bacteria, viruses, fungi, and protozoa, sourced from the NCBI RefSeq database on January 22, 2023 (kraken2-build --build). Before constructing the database, we masked repetitive regions in reference genomes using Dustmasker to prevent false positives in the results [7]. Subsequently, we applied Bracken (v2.7.0) to further refine the outcomes [8]. For this, we constructed Bracken databases using the options (-k 35 -l 150). We set a taxa threshold of minimum 10 reads to ensure sufficient confidence. To evaluate Kraken's classification accuracy and differentiate between reads that are distributed throughout a reference genome and those associated with only a small segment (potentially indicating false positives), we used KrakenUniq to estimate the number of distinct k-mers linked to each taxon in the input sequencing data [9]. This feature was integrated into Kraken2 by enabling the relevant options (--report-minimizer-data).

**Statistical analysis**

We computed the richness (observed number of features) and Shannon–Wiener diversity index, which assesses the uncertainty related to predicting the identity of a randomly chosen individual within the community, for both taxonomic and functional features using the R package 'vegan'. To explore the connections between composition, richness, diversity of taxonomic and functional features, and environmental factors (such as daily mean $PM_{10}$ concentration, temperature, and relative humidity), we conducted Pearson's correlation analysis. This analysis was performed using the R package 'Hmisc'.

Principal Coordinates Analysis (PCoA) was used as the ordination method based on Bray–Curtis dissimilarity of taxonomic and functional compositional data (i.e., relative taxa or functional feature abundance, calculated as the reads assigned to a taxa or function divided by the sum), and this analysis was conducted using the R package 'vegan'. The heatmap representing the Bray–Curtis dissimilarity of taxonomic and functional compositional data was visualized using the R package 'ComplexHeatmap'.

We performed an ANOVA analysis using distance matrices to assess the extent to which the variation in the composition of the atmospheric microbiome (i.e., both taxonomic and functional features) could be attributed to individual variables. For this analysis, we employed the Adonis2 function from the 'vegan' package, which performs permutational MANOVA, utilizing 1000 permutations and the Bray–Curtis dissimilarity matrix of taxonomic and functional compositional data (i.e., relative abundance). The following formula design was employed: daily mean $PM_{10}$ + air mass origin + daily mean relative humidity + daily mean temperature, in the specified order. Significance was determined based on a threshold of P-value below 0.05. Furthermore, we assessed the differences in the composition of the atmospheric microbiome between dusty and clear atmospheric conditions using the same methodology. To evaluate the multivariate homogeneity of group dispersions, an assumption of MANOVA, we employed the Betadisper function with 1000 permutations.

In all results, including both read and assembly-based approaches, we added a pseudocount of 1 to the counts (CPM and RPKM) before log transformation, if necessary. We applied a filtering criterion to focus our analysis on taxonomic and functional features with a significant presence in the dataset. Specifically, only features with a positive read number in at least 1/4 of all samples (providing a sample size of more than 12 observations per taxon or functional feature) were included for both correlation and differential relative abundance analysis. This criterion prioritizes features that exhibited consistent presence throughout the dataset, ensuring the robustness and reliability of the analyses. In the correlation analysis between contigs associated with antibiotic resistance, virulence, and pathogenicity, and daily mean $PM_{10}$ and temperature, we excluded the air samples collected on May 19, 2022, due to concerns regarding potential high anthropogenic contributions on that specific day, likely attributed to a holiday celebrated with large bonfires throughout the country on the night before the sampling occurred.

We used the Shapiro–Wilk normality test to assess data distribution normality. The test results indicated a failure to meet this assumption. Therefore, we chose a nonparametric method for comparing the two groups. The Wilcoxon rank sum test, also known as the Mann–Whitney *U* test, was utilized to assess the differential relative abundance of taxa between the dusty and clear atmospheric conditions. To account for multiple comparisons, we applied the Benjamini–Hochberg correction to control for false discovery rate, with the adjusted P-value below 0.05 indicating statistical significance.

We assessed the relationship between specific microbial features (abundance of species and contigs associated with antibiotic resistance, virulence, and pathogenicity) and environmental variables (including daily mean $PM_{10}$ concentration and temperature) using Pearson’s correlation. A statistically significant correlation was considered for an absolute Pearson's correlation coefficient ($r$) value above 0.3 and a P-value below 0.05.

In our study, we chose Pearson’s correlation to assess the relationship between specific microbial features and environmental variables, including daily mean $PM_{10}$ concentration and temperature. This choice was motivated by several considerations. Firstly, Pearson's correlation is well-suited when linear associations are of particular interest, which aligns with our preliminary results showing linear trends between microbial features and environmental variables. Secondly, the absence of outliers was a key factor. Although our data includes extreme events like dust storms, where particulate matter concentrations are inherently elevated, these values and their corresponding effects on microbial features are integral to our investigation rather than outliers. However, to mitigate any potential exaggerated effects caused by extreme values on specific days, we applied a $\log_{10}$ transformation to the abundance of microbial features and daily $PM_{10}$ and temperature values in these correlation analyses. Additionally, each correlation analysis between microbial feature abundance and environmental factors represents an independent hypothesis test, without simultaneous comparisons across different taxa or features. Therefore, we chose not to include multiple testing corrected P-values in our results unless otherwise mentioned. This decision acknowledges that stringent corrections might be overly conservative for independent tests in our analysis. Multiple correction methods, such as Bonferroni or False Discovery Rate, are typically recommended when conducting many simultaneous tests (such as the co-occurrence analysis of microbial species in this study) to control the increased risk of Type I errors. However, in our case, each correlation represents a distinct analysis, and the independent nature of these comparisons reduces the likelihood of inflating the overall Type I error rate.

While the observed linear associations in our preliminary analysis indicated that using Pearson's correlation would be a better fit for the correlation analysis among the microbial species, for clarity, we also conducted the same analysis using Spearman's correlation. Spearman is a non-parametric correlation test known for assessing the monotonic relationship between variables. The results of this analysis showed that, although similar trends to Pearson's correlation were observed in Spearman's correlation, as shown in Supporting Figure 24, an inflated number of significant correlations were observed in Spearman, supporting our decision to use Pearson's correlation.

We conducted a co-occurrence analysis of microbial species based on Pearson's correlation to explore the relationship between specific taxa. Only results meeting the following criteria were considered statistically robust between taxa and retained for further examination: an absolute Pearson's correlation coefficient ($r$) value above 0.3 and a P-value below 0.05, after adjustment by Benjamini–Hochberg's method to reduce the chances of obtaining false-positive results.

#### Results

##### DNA Yield of operational blanks and contamination assessment

The DNA yield from the operational blanks, which serve as control samples to assess background contamination, was either close to the detection limit (0.005 ng/μL) or lower. As a result, both shotgun metagenome sequencing (with only three out of seven samples generating sequencing results) and Kraken2 taxonomic classification of these control samples displayed substantially lower read counts compared to air samples. This strongly supports a minimal likelihood of background contamination (details can be found in Supporting Tables 1, 2 and 13).

##### Meteorological conditions before and during the sampling period and, the estimated origin of dust

We collected air samples from March 29$^{th}$ to May 31$^{st}$, 2022, at the onset of the dry season after winter rainfalls. We did not record any precipitation during this period (Supporting Figure 2). Temperatures steadily increased during the campaign, ranging from 16.5 to 27.1°C (Supporting Figure 3). We defined dust events using a daily mean $PM_{10}$ concentration threshold of 65.2 μg $m^{-3}$, based on a recent study conducted in the EM [10]. Over the sampling period, only fifteen days exceeded this threshold, indicating dusty conditions. We identified three distinct air transport patterns using dust column mass density maps and back trajectory analysis of air masses: northwesterly, southwesterly, and easterly (Supporting Movies 1-4). In summary, we collected a total of forty-five air samples during the sampling period, comprising both dusty (n=13) and clear (n=32) atmospheric conditions (Supporting Figure 3 and Supporting Table 3).

### Figures

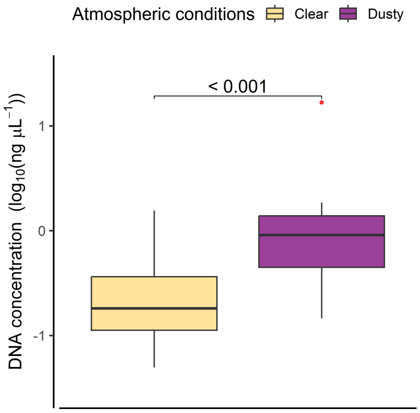


**Supporting Figure 1.** DNA yield from air samples collected during different atmospheric conditions. The statistical significance between dusty and clear atmospheric conditions was assessed using the Mann–Whitney *U* test. Boxplot center lines represent the median values, while the lower and upper hinges correspond to the first and third quartiles ($25^{th}$ and $75^{th}$ percentiles, respectively). Whiskers extend from the hinges to the lowest and largest values within 1.5 times the Interquartile Range (IQR) from the hinge, with outliers depicted as red-colored dots.

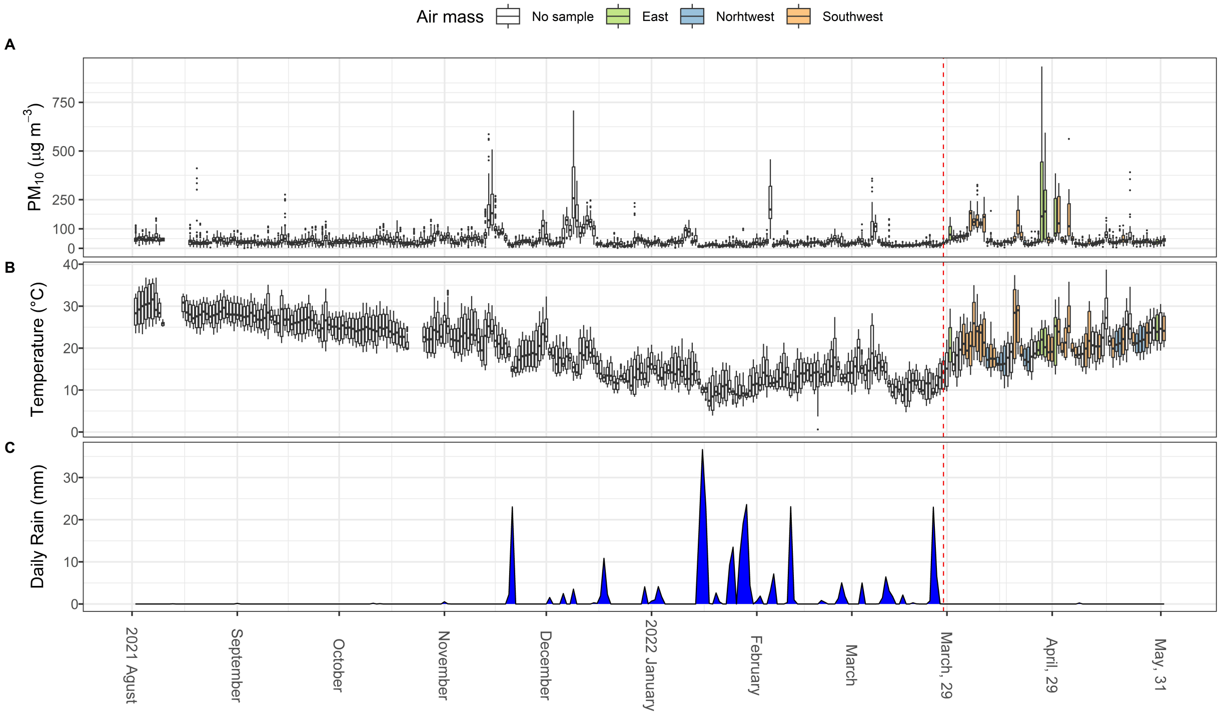


**Supporting Figure 2.** Meteorological conditions before and during the sampling period. Panel (A) showcases the daily distribution of particulate matter concentration ($PM_{10}$), while panel (B) illustrates the corresponding temperature variations. Additionally, panel (C) portrays the daily accumulation of rainfall. In panels A and B, the center lines of the boxplots represent the median values, while the lower and upper hinges indicate the first and third quartiles ($25^{th}$ and $75^{th}$ percentiles, respectively). The lower and upper whiskers extend from the hinges to the lowest and largest values within 1.5 times the Interquartile Range (IQR) from the hinge. Outliers are denoted as red-colored dots. Red dotted lines signify the commencement of the sampling campaign. Within the sampling period, white-colored box plots labeled as 'No sample' indicate days when air sampling was omitted.

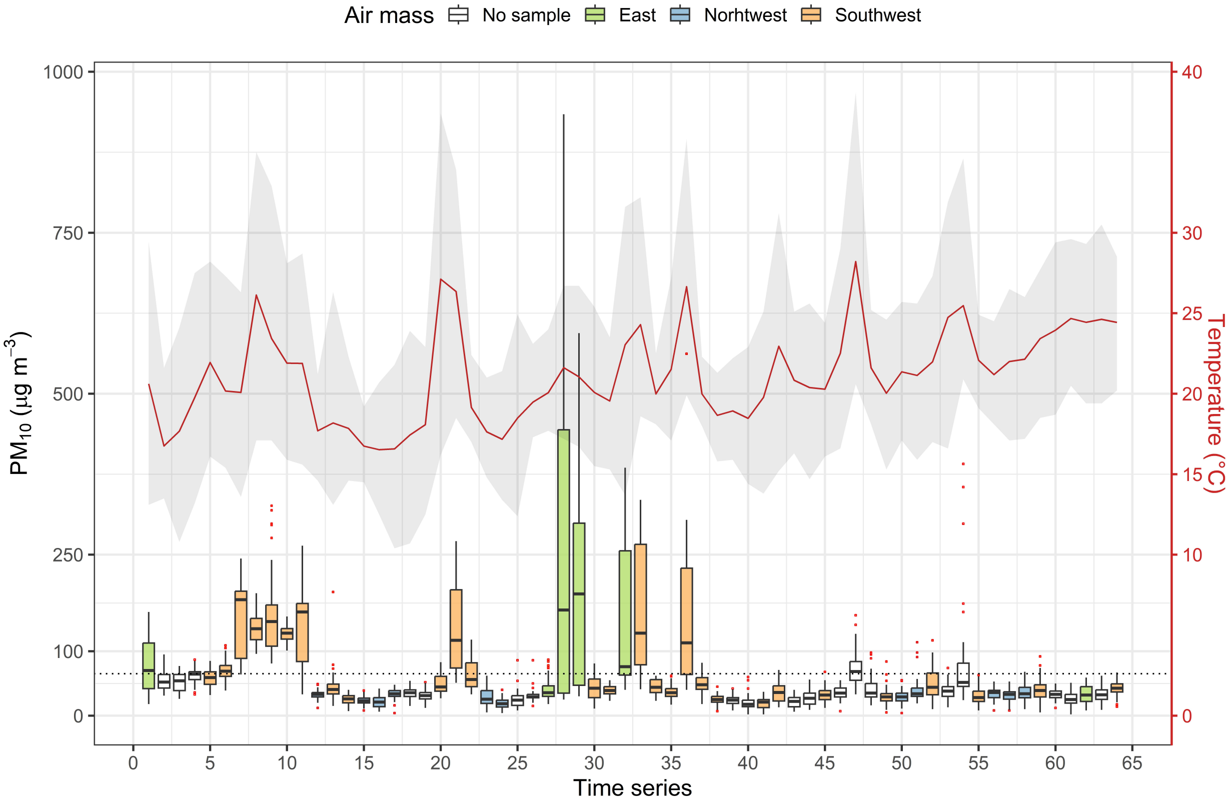


**Supporting Figure 3.** Meteorological conditions during the sampling period (March 29th to May 31st, 2022). The x-axis displays the distribution of daily $PM_{10}$ concentrations, while the y-axis represents the variations in daily temperature. The red line indicates the daily mean temperature, and the gray-colored area illustrates the range between the daily minimum and maximum temperature values. The color coding in the figure represents the air mass origins for the respective data points. Center lines in the boxplots indicate the median values, with lower and upper hinges representing the first and third quartiles (25th and 75th percentiles, respectively). Whiskers extend from the hinges to the lowest and largest values within 1.5 times the Interquartile Range (IQR) from the hinge, while outliers are depicted as red-colored dots. During the sampling campaign, white-colored box plots labeled as 'No sample' denote days when air sampling was omitted. A dotted black line within the plot signifies dust events, defined by a $PM_{10}$ concentration threshold of 65.2 µg $m^{-3}$, established based on a recent study conducted in the EM, as detailed in the methods section of the main manuscript.

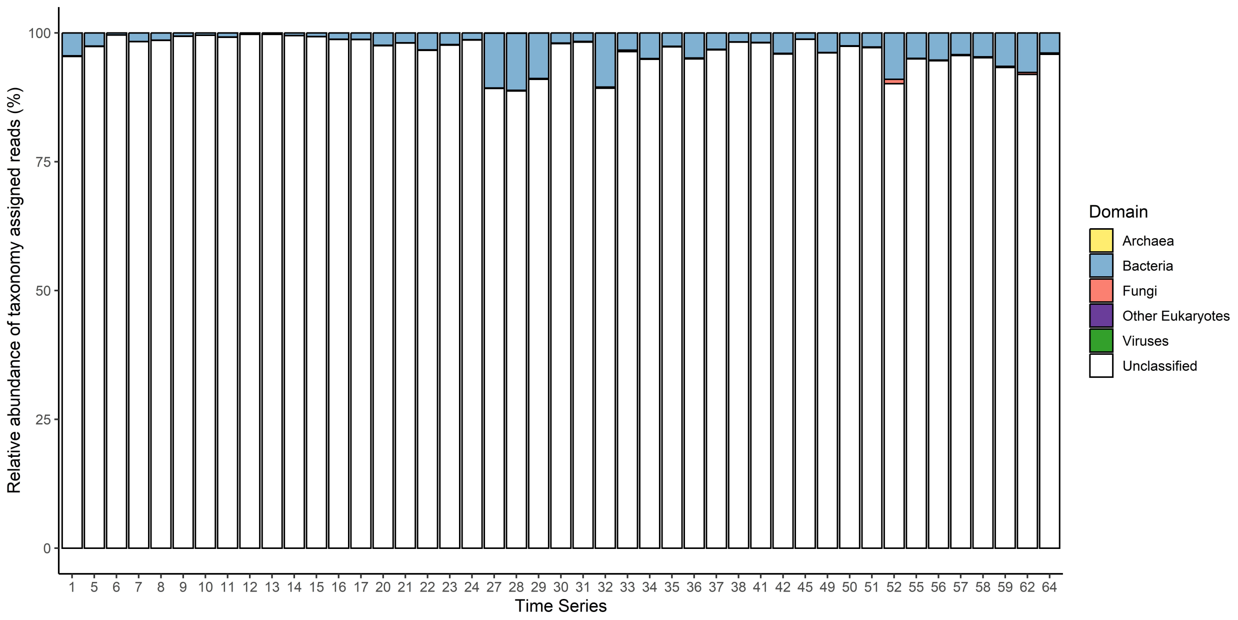


**Supporting Figure 4.** Relative abundance of taxonomy-assigned reads (%). The category "Unclassified" denotes sequences for which a label at the root of the taxonomic tree did not find a matching k-mer in any reference genomes or exceed the specified threshold (--confidence 0.1) in Kraken2, resulting in them being labeled as unclassified.

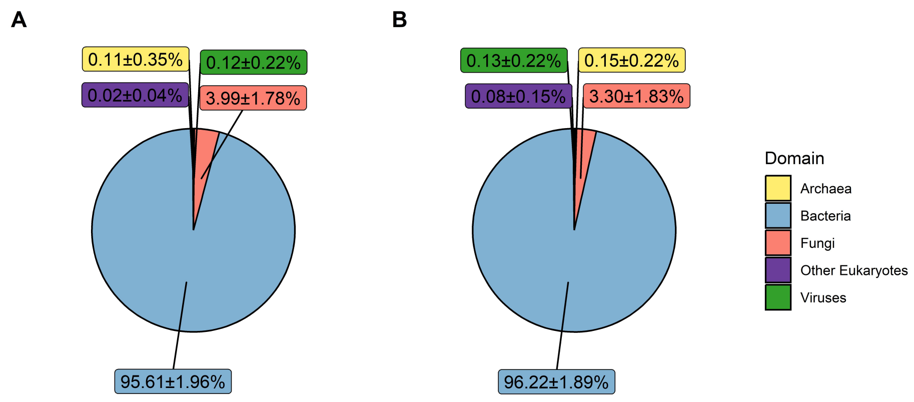


**Supporting Figure 5.** Relative abundances of metagenomic reads categorized into taxonomic groups in clear (A) and dusty (B) atmospheric conditions. Values within their respective domains are presented as the mean relative abundance ± standard deviation (SD) (%).

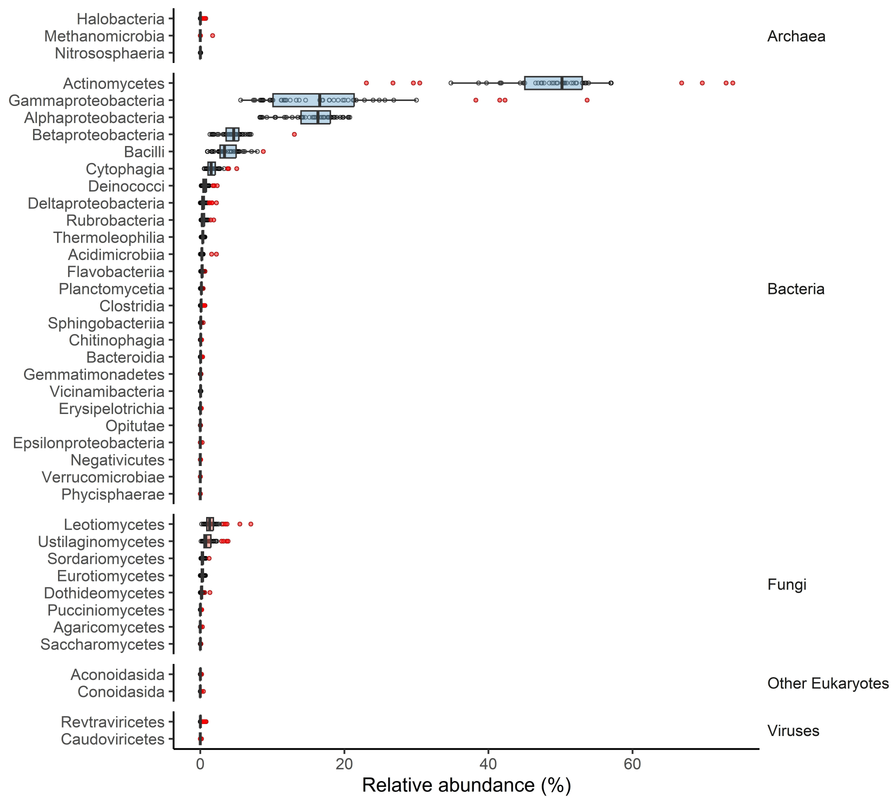


**Supporting Figure 6.** Distribution of microbial relative abundance (%) at the class level observed in at least one-quarter of total samples. Boxplot center lines depict the median values, while the lower and upper hinges represent the first and third quartiles (25$^{th}$ and 75$^{th}$ percentiles). Whiskers extend from the hinges to the lowest and largest values within 1.5 times the Interquartile Range (IQR) from the hinge. Individual samples within each group are shown as dots, with outliers presented as red-colored dots.

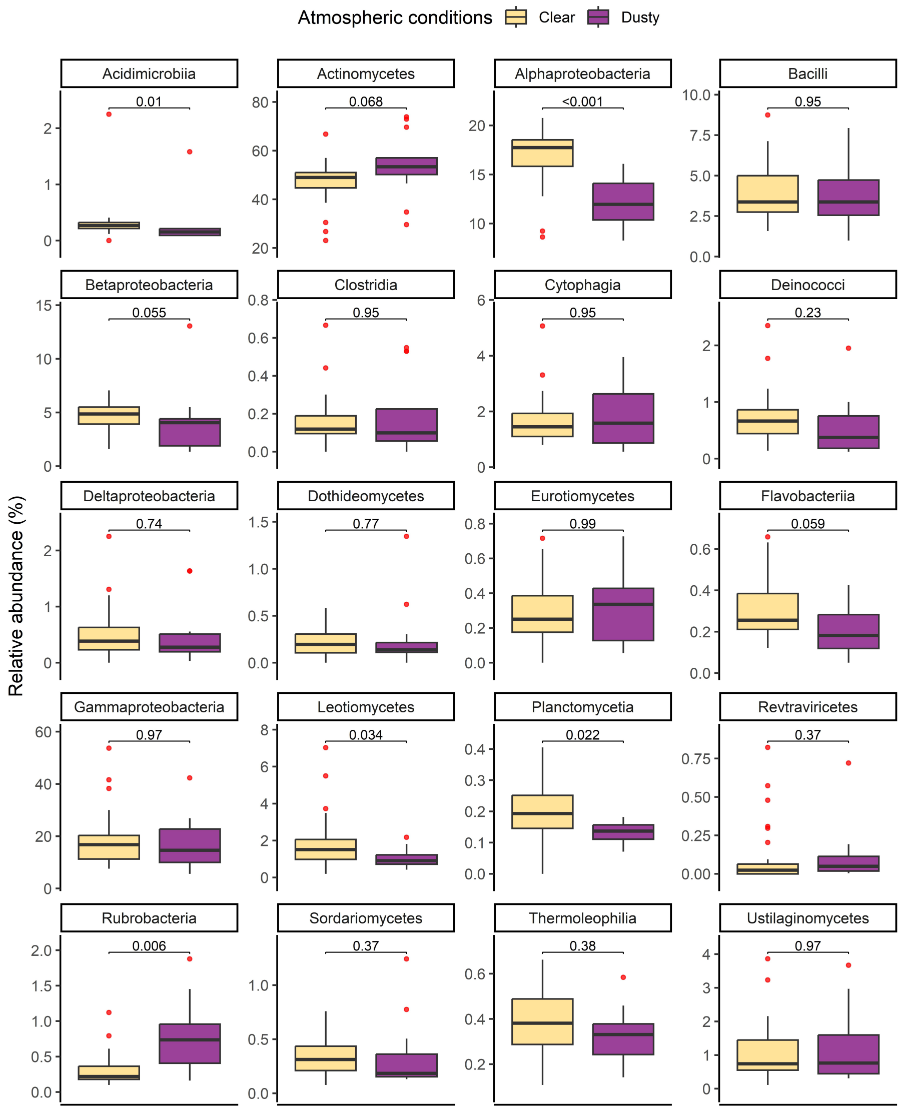


**Supporting Figure 7.** Relative abundance of significant microbial classes observed in at least one-fourth of all samples, comparing dusty and clear atmospheric conditions. Statistical significance between these conditions was evaluated using the Mann–Whitney *U* test, and P-values were adjusted using the Benjamini–Hochberg method to reduce false-positive outcomes. The figure illustrates the most abundant 20 taxa. Boxplot center lines represent median values, while the lower and upper hinges correspond to the first and third quartiles (25$^{th}$ and 75$^{th}$ percentiles). Whiskers extend from the hinges to the lowest and largest values within 1.5 times the Interquartile Range (IQR) from the hinge. Red-colored dots denote outliers.

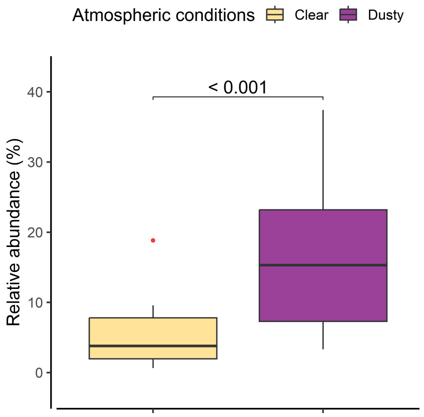


**Supporting Figure 8.** Relative abundance (%) of *G. obscurus*, a taxon of specific interest within Actinomycetes, observed in dusty and clear atmospheric conditions. Statistical significance between these conditions was assessed using the Mann–Whitney *U* test across all species observed in at least one-fourth of all samples. Subsequently, P-values underwent adjustment using the Benjamini–Hochberg method to mitigate the risk of false-positive outcomes. Boxplot center lines represent median values, while the lower and upper hinges correspond to the first and third quartiles ($25^{th}$ and $75^{th}$ percentiles). Whiskers extend from the hinges to the lowest and largest values within 1.5 times the Interquartile Range (IQR) from the hinge. Red-colored dots denote outliers.

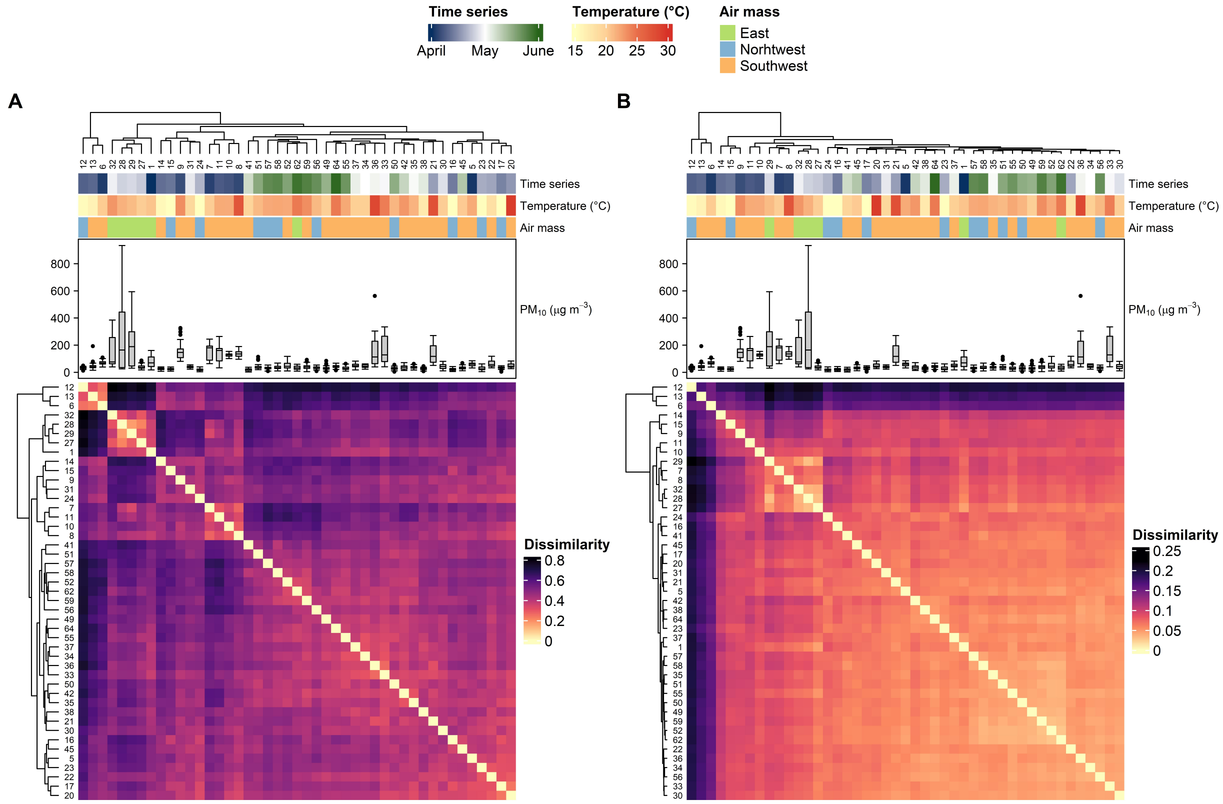


**Supporting Figure 9.** Heatmap illustrating Bray–Curtis dissimilarity of (A) taxonomic and (B) functional compositional data across the time series. The panels represent each day of the time series, starting from day 1 on March 29th to day 64 on May 31st, 2022. The heatmap also includes daily mean temperature, air mass origin, and distribution of daily mean $PM_{10}$ concentration. Boxplot center lines depict the median values, while the lower and upper hinges represent the first and third quartiles (25th and 75th percentiles). Whiskers extend from the hinges to the lowest and largest values within 1.5 times the Interquartile Range (IQR) from the hinge. Outliers are displayed as black-colored dots.

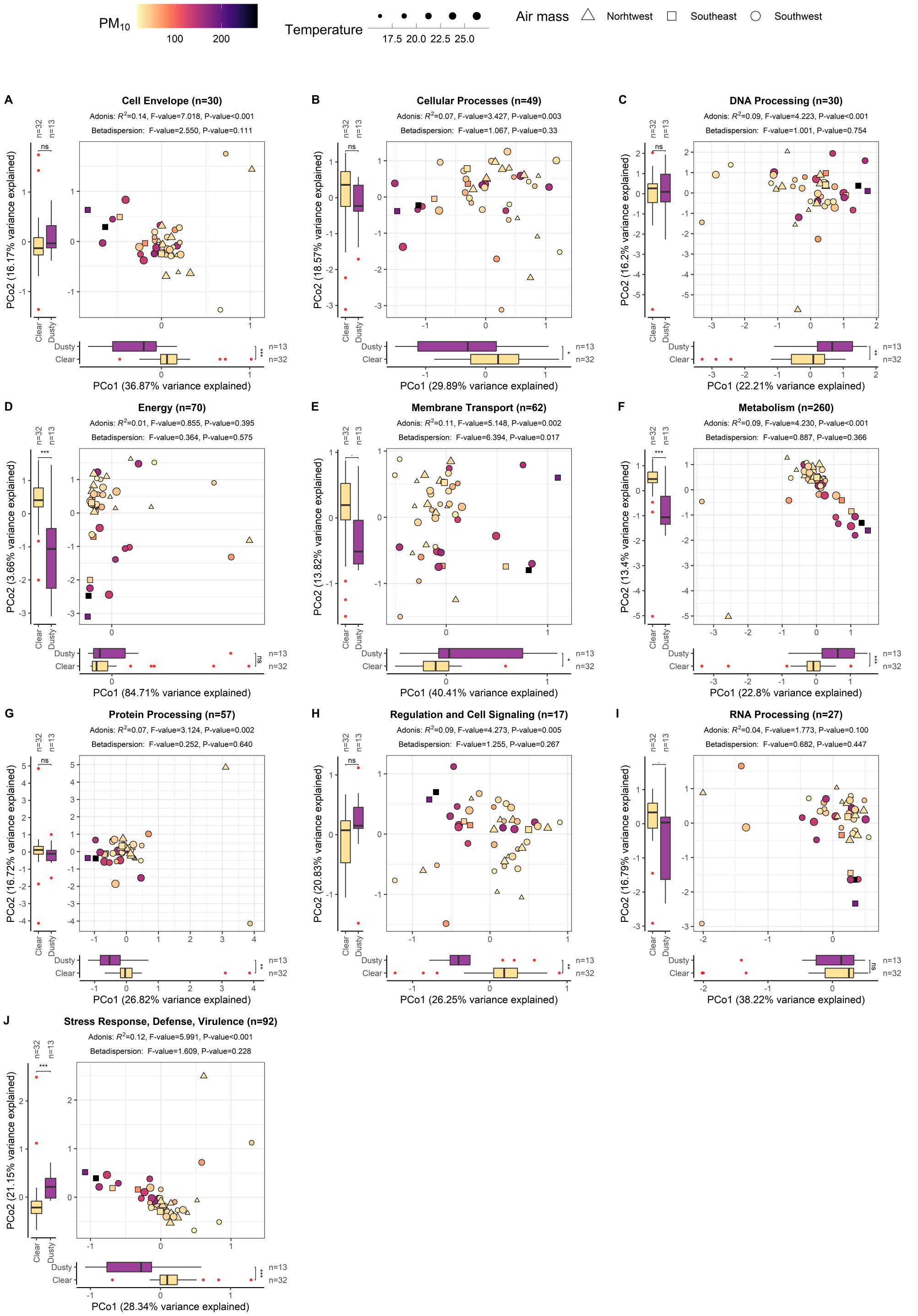


**Supporting Figure 10.** Functional composition across atmospheric conditions for specific subcategories (A-J) within the atmospheric microbiome. Each plot denotes a functional category illustrating variations in $PM_{10}$ concentration, temperature, and air mass origin. 'n' indicates the count of functional features in each subcategory. Principal Coordinate Analysis (PCoA) plots demonstrate the proportion of variance explained by each axis. Adonis test results, based on Bray–Curtis dissimilarity, are presented below each plot's title. Multivariate homogeneity of group dispersions (betadispersion), a prerequisite of permutational MANOVA, is also displayed. Mann–Whitney *U* test assessed significant differences in PCoA axes between dusty and clear conditions. Boxplots display median values (center lines), first and third quartiles (lower and upper hinges, 25th and 75th percentiles), while whiskers extend from hinges to values within 1.5 times the Interquartile Range (IQR). Outliers are red dots. Significance levels are indicated by symbols: '***' for <0.001, '**' for <0.01, '*' for <0.05, '.' for <0.1, and 'ns' for <1.

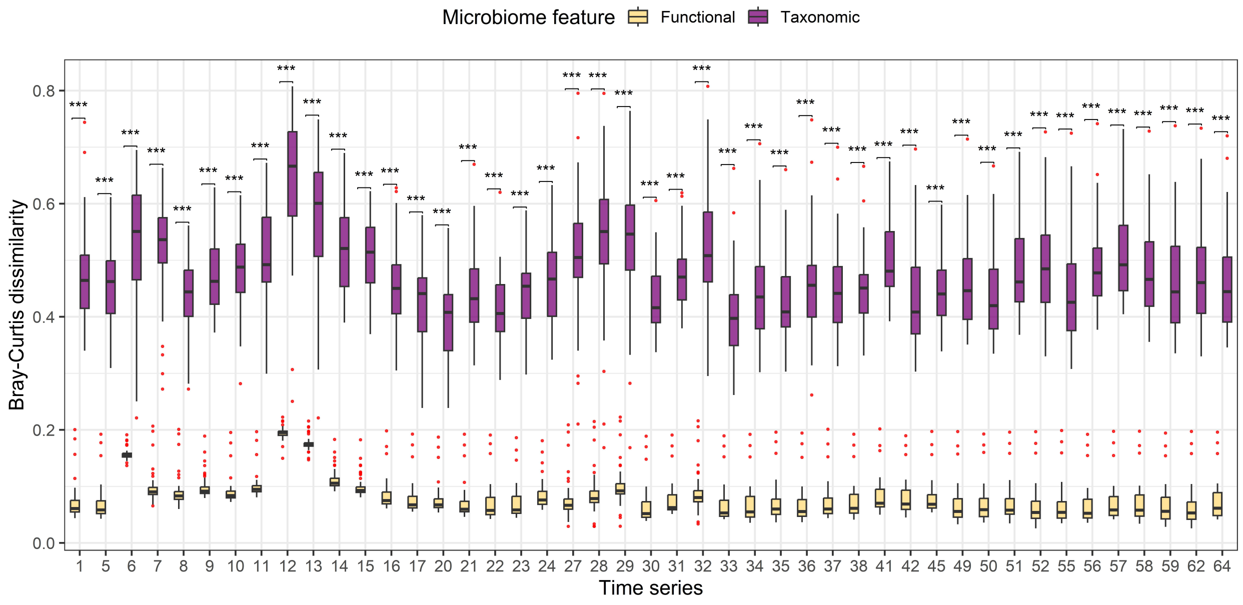


**Supporting Figure 11.** Comparison of Bray–Curtis dissimilarity in taxonomic and functional features for each sample over a time series. The initial sample (day 1) was collected on March 29$^{th}$, and the final sample (day 64) was obtained on May 31$^{st}$, 2022. Statistical significance between these conditions was assessed using the Mann–Whitney *U* test across all samples. Subsequently, P-values underwent adjustment using the Benjamini–Hochberg method to minimize the risk of false-positive outcomes. Boxplot center lines represent the median values, while the lower and upper hinges indicate the first and third quartiles (25$^{th}$ and 75$^{th}$ percentiles). Whiskers extend from the hinges to the lowest and largest values within 1.5 times the Interquartile Range (IQR) from the hinge. Outliers are shown as red-colored dots. Significance levels are indicated by symbols: '***' for <0.001, '**' for <0.01, '*' for <0.05, '.' for <0.1, and 'ns' for <1.

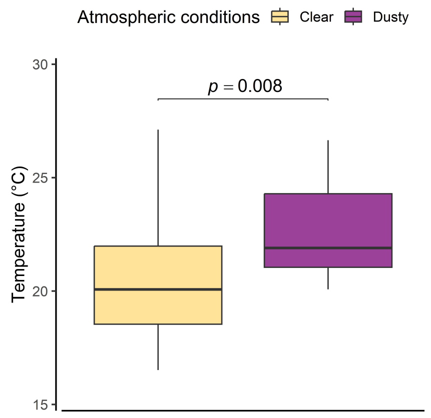


**Supporting Figure 12.** Daily mean temperature values during dusty and clear atmospheric conditions. The statistical significance between these conditions was assessed using the Mann–Whitney *U* test. Boxplot center lines represent the median values, while the lower and upper hinges correspond to the first and third quartiles (25$^{th}$ and 75$^{th}$ percentiles). The lower and upper whiskers extend from the hinges to the lowest and largest values within 1.5 times the Interquartile Range (IQR) from the hinge. Outliers are displayed as red-colored dots.

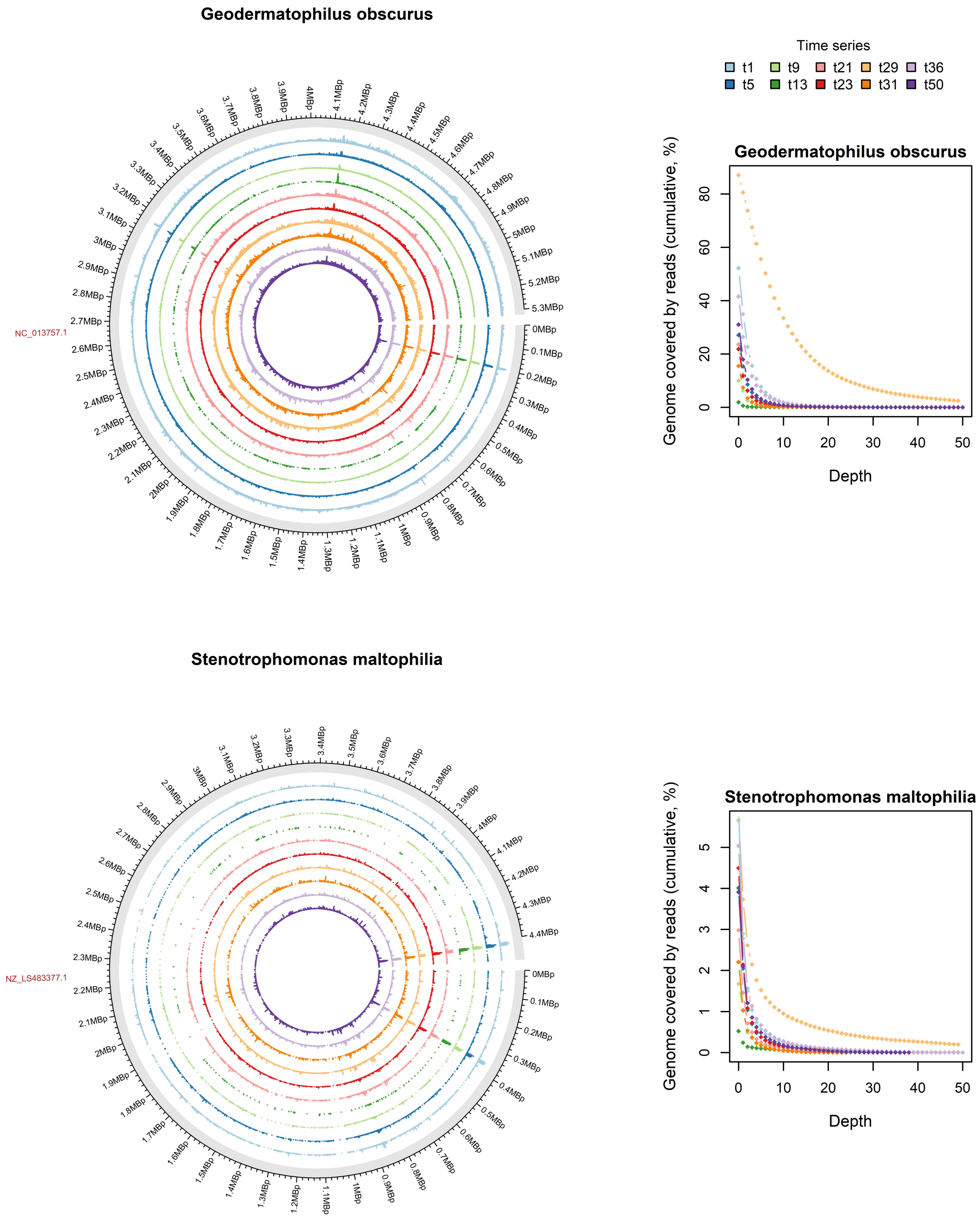


**Supporting Figure 13.** Read mapping results of individual samples to reference genomes, demonstrating coverage and depth. The circus plots show the read mapping results for the most prevalent taxa across all samples, with *G. obscurus* serving as the reference, accompanied by the predominant bacterial pathogens *S. maltophilia*. The outer circle represents the entire genome of each species, including chromosomes and plasmids, with a gap denoting the boundary. The RefSeq number is displayed at the top, outside the Circos plot, and genome size is annotated by numbers around each circle. Inside, colored circles (bar plots) depict air samples collected under various atmospheric conditions. For visual convenience, only five clear days (t5, t13, t23, t31 and t50) and five dusty days (t1, t9, t21, t29 and t36) are shown here. Each aligned read within a sample to the reference genome is depicted as a colored bar, indicating its specific position in the reference genome, with the height of the bar representing the depth. Coverage histograms illustrate the cumulative percentage of the total number of bases in the reference genome covered by different read depths.

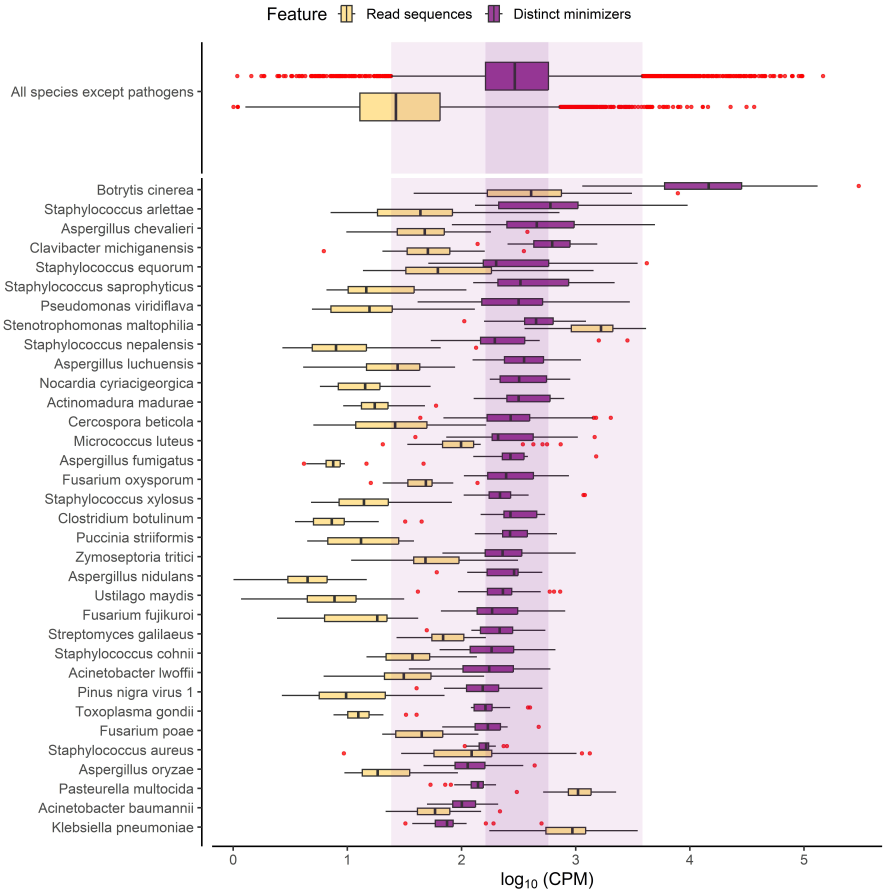


**Supporting Figure 14.** A comparative analysis between Kraken2's read abundance and KrakenUniq's count of distinct minimizers attributed to each taxon within the input sequencing data. The comparison specifically focuses on selected taxa of interest in relation to their distribution across all samples for all taxa. Boxplot visualizations display key statistical characteristics: center lines represent the median values, while the lower and upper hinges delineate the first and third quartiles ($25^{th}$ and $75^{th}$ percentiles, respectively). Whiskers extend from the hinges to the lowest and largest values within 1.5 times the Interquartile Range (IQR) from the hinge. Red-colored dots represent outliers. Additionally, the dark purple shade denotes the minimum and maximum values within the boxplot for KrakenUniq's count of distinct minimizers, while the light shade encapsulates the lower and upper hinges of the boxplot, representing the range for KrakenUniq's distinct minimizer count across all species. This comparison aids in assessing the relationship between read abundance and distinct minimizer counts for specific taxa of interest.

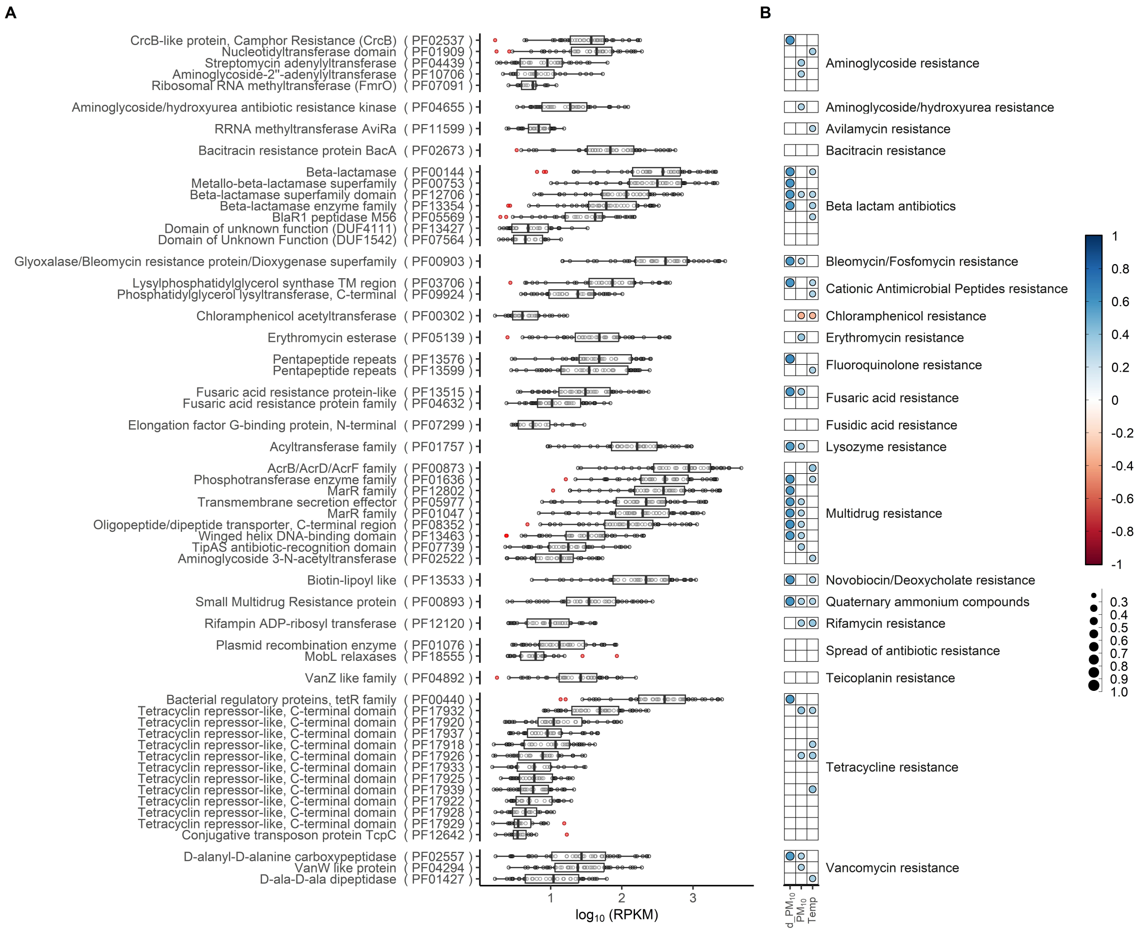


**Supporting Figure 15.** The relationship between functional features associated with antibiotic resistance and environmental variables. (A) Abundance distribution ($\log_{10}$(RPKM)) for the most prevalent PFAM entries associated with antibiotic resistance. Boxplot details: center lines depict medians, hinges represent the 25$^{th}$ and 75$^{th}$ percentiles, while whiskers extend to values within 1.5 times the Interquartile Range (IQR) from the hinges. Red-colored dots indicate outliers. (B) Pearson's correlation analysis between individual taxa and environmental variables. 'd_$PM_{10}$' represents the correlation between daily mean $PM_{10}$ concentration specifically within dusty atmospheric conditions, while '$PM_{10}$' denotes the correlation across all samples. 'Temp' signifies the correlation between each taxon and the daily mean temperature. The visual representation using color-coded Pearson's correlation coefficient (*r*) values and different-sized bins assists in interpreting the data visually. PFAM entries are categorized based on their resistance to antibiotic drug classes for improved clarity.

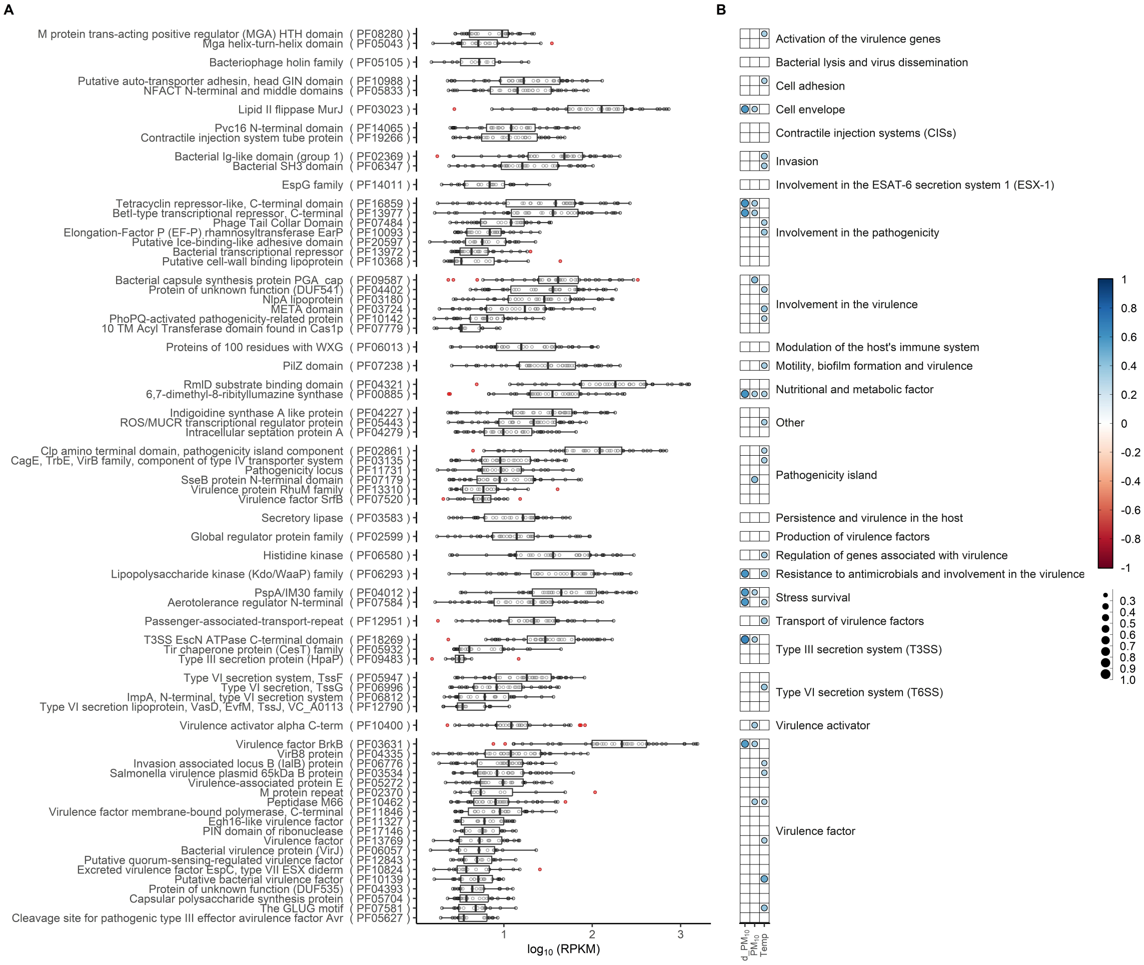


**Supporting Figure 16.** The relationship between functional features associated with virulence and pathogenicity and environmental variables. (A) Abundance distribution ($\log_{10}$(RPKM)) for the most prevalent PFAM entries associated with virulence and pathogenicity. Each PFAM entry's accession number is presented following the name in parentheses. Boxplot details: center lines represent median values, while lower and upper hinges depict the 25$^{th}$ and 75$^{th}$ percentiles, respectively. The lower and upper whiskers extend from the hinges to the lowest and largest values within 1.5 times the Interquartile Range (IQR) from the hinge. Outliers are shown as red-colored dots. (B) Pearson's correlation analysis between individual taxa and environmental variables. 'd_$PM_{10}$' signifies the correlation between daily mean $PM_{10}$ concentration specifically within dusty atmospheric conditions, while '$PM_{10}$' denotes the correlation across all samples. 'Temp' represents the correlation analysis between each taxon and the daily mean temperature. The visual representation using color-coded Pearson's correlation coefficient ($r$) values and different-sized bins aids in data interpretation. PFAM entries are categorized based on their virulence and pathogenicity mechanisms for clarity.

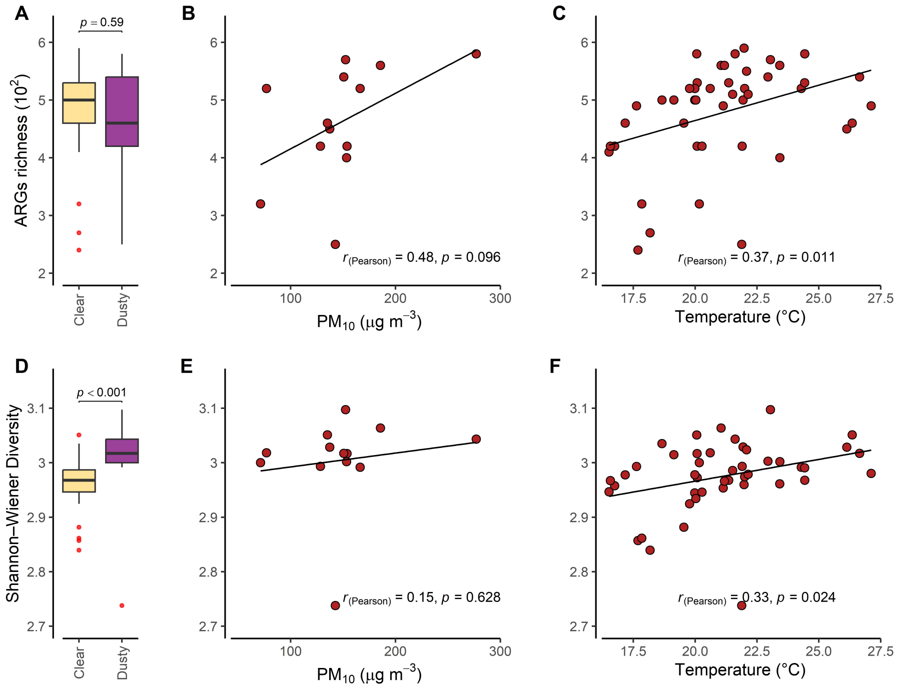


**Supporting Figure 17.** Richness and diversity of antibiotic resistance-contigs. (A) Richness and (D) diversity plots illustrate the richness and diversity of antibiotic resistance-contigs under dusty and clear conditions. Pearson's correlation analysis explores the relationship between (B, E) $PM_{10}$ concentration (exclusive to dusty atmospheric conditions), (C, F) temperature across all samples, and microbiome richness/diversity indices. Boxplots display median values (center lines), first and third quartiles (lower and upper hinges, $25^{th}$ and $75^{th}$ percentiles), while whiskers extend from hinges to values within 1.5 times the Interquartile Range (IQR). Outliers are represented as red dots. The Mann–Whitney *U* test assessed significant differences in richness and diversity between dusty and clear conditions.

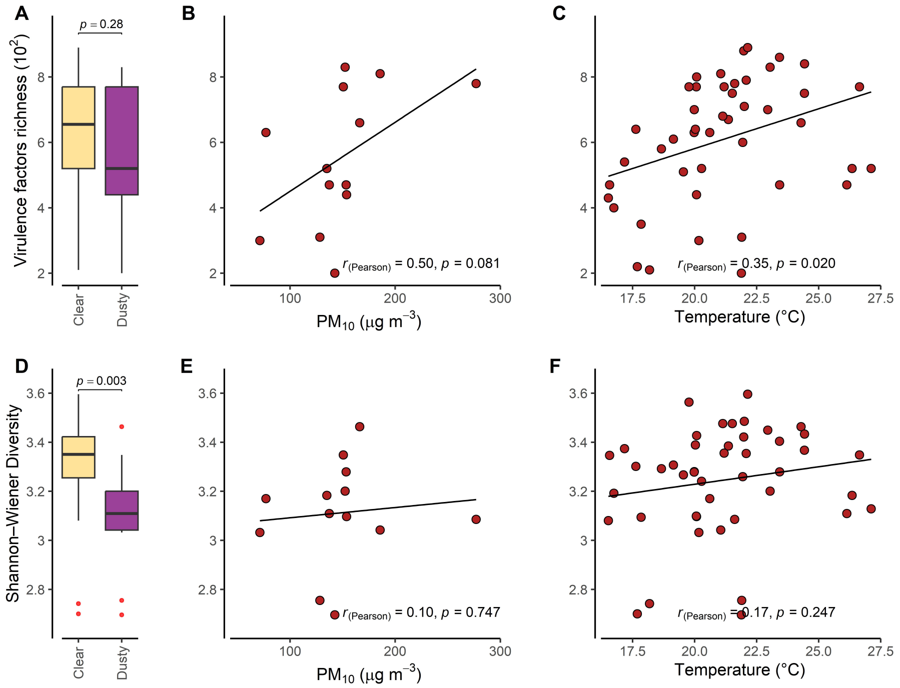


**Supporting Figure 18.** Richness and diversity of virulence-related contigs. (A) Richness and (D) diversity plots illustrate the richness and diversity of virulence-related-contigs under dusty and clear conditions. Pearson's correlation analysis explores the relationship between (B, E) $PM_{10}$ concentration (exclusive to dusty atmospheric conditions), (C, F) temperature across all samples, and microbiome richness/diversity indices. Boxplots display median values (center lines), first and third quartiles (lower and upper hinges, $25^{th}$ and $75^{th}$ percentiles), while whiskers extend from hinges to values within 1.5 times the Interquartile Range (IQR). Outliers are represented as red dots. The Mann–Whitney *U* test assessed significant differences in richness and diversity between dusty and clear conditions.

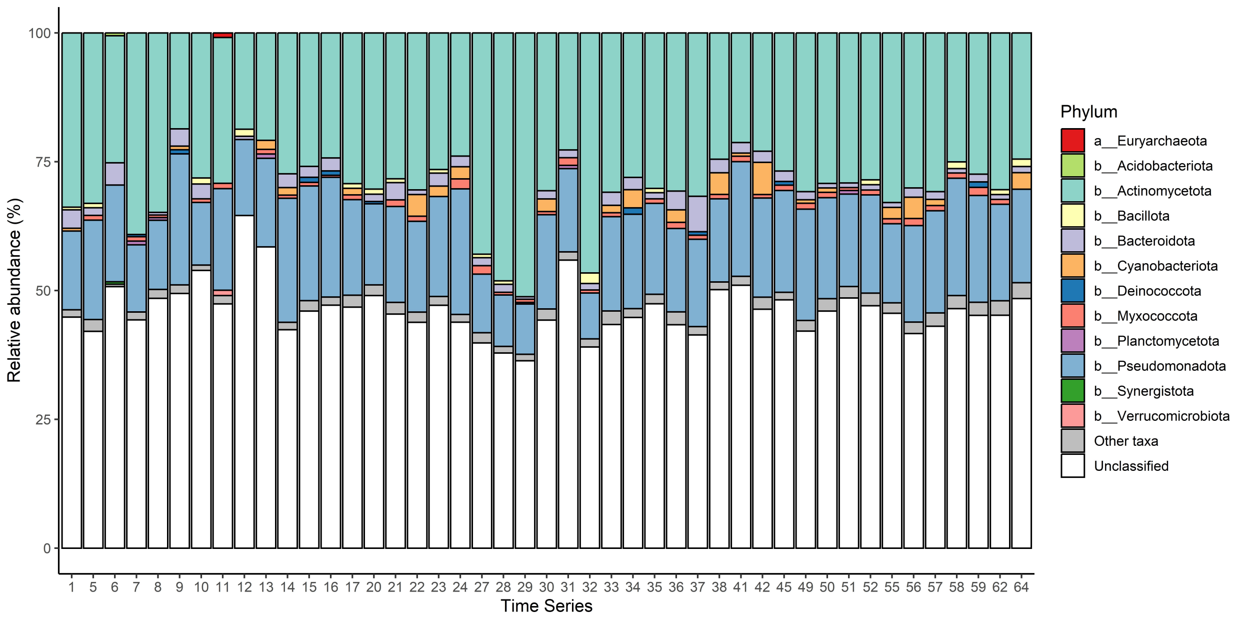


**Supporting Figure 19.** Relative abundance of host microorganisms carrying antibiotic resistance traits (%) within the dataset. This figure specifically represents contigs associated with antibiotic resistance traits. The category "Unclassified" indicates antibiotic resistance-contigs for which a label at the root of the taxonomic tree did not find a matching k-mer in any reference genomes in Kraken2, hence labeled as unclassified. In the figure legend, 'a', 'b', and 'e' signify Bacteria, Archaea, and Eukarya, respectively.

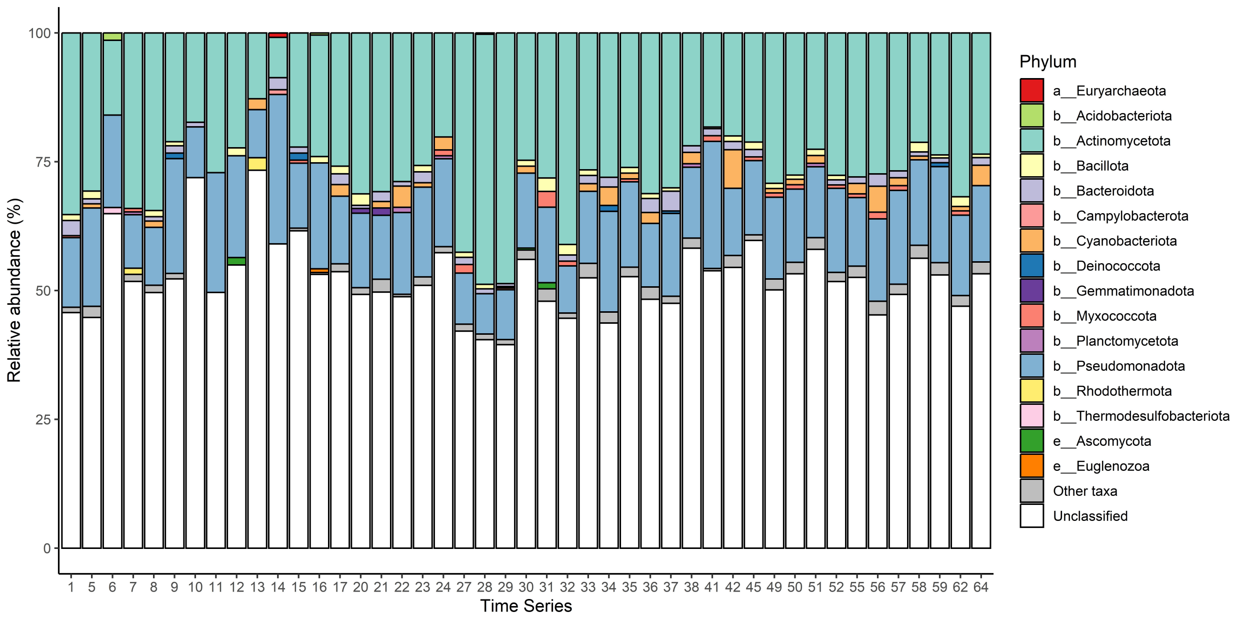


**Supporting Figure 20.** Relative abundance of host microorganisms carrying virulence-related traits (%) within the dataset. This figure specifically represents contigs associated with virulence-related traits. The category "Unclassified" indicates virulence-related-contigs for which a label at the root of the taxonomic tree did not find a matching k-mer in any reference genomes in Kraken2, hence labeled as unclassified. In the figure legend, 'a', 'b', and 'e' signify Bacteria, Archaea, and Eukarya, respectively.

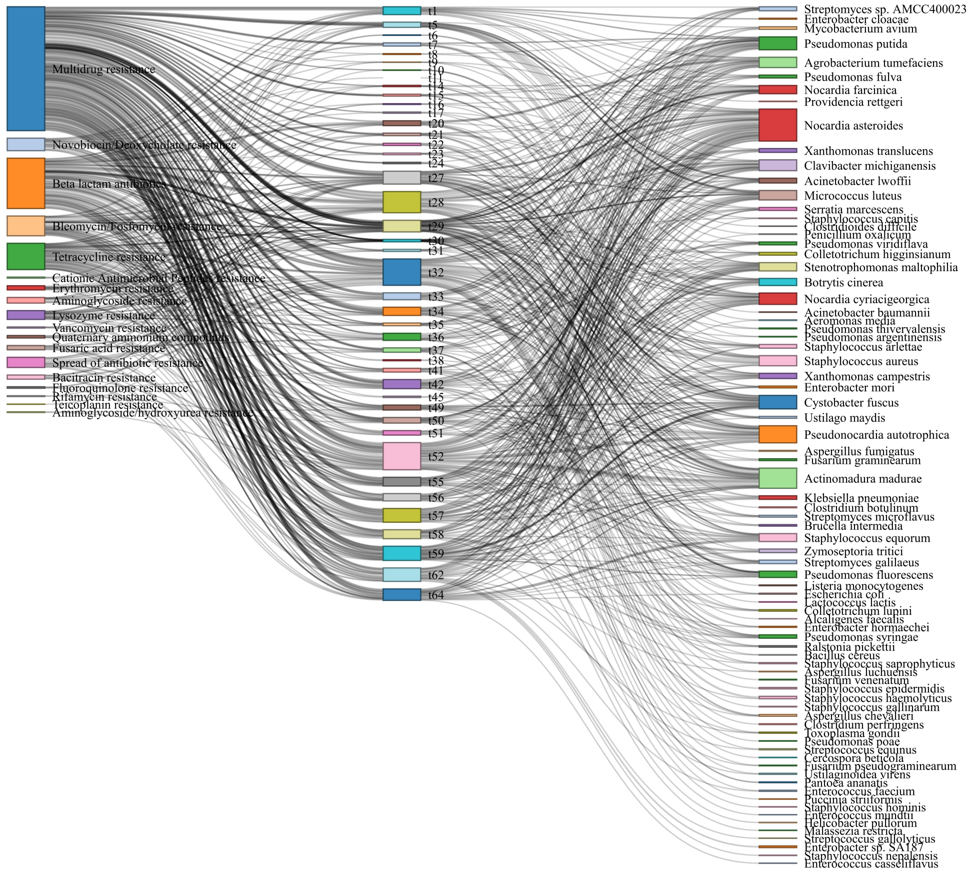


**Supporting Figure 21.** Host microorganisms carrying antibiotic resistance traits with known pathogenicity to humans, animals, and plants found in air samples. The length of the rectangles indicates the RPKM values. From left to right, the rectangles represent antibiotic resistance traits, sampling day (time series), and potential pathogens. For visual convenience, RPKM values were not depicted, but for reference, the total abundance of Multidrug resistance-genes (blue colored rectangle left-top) is 791.95 RPKM.

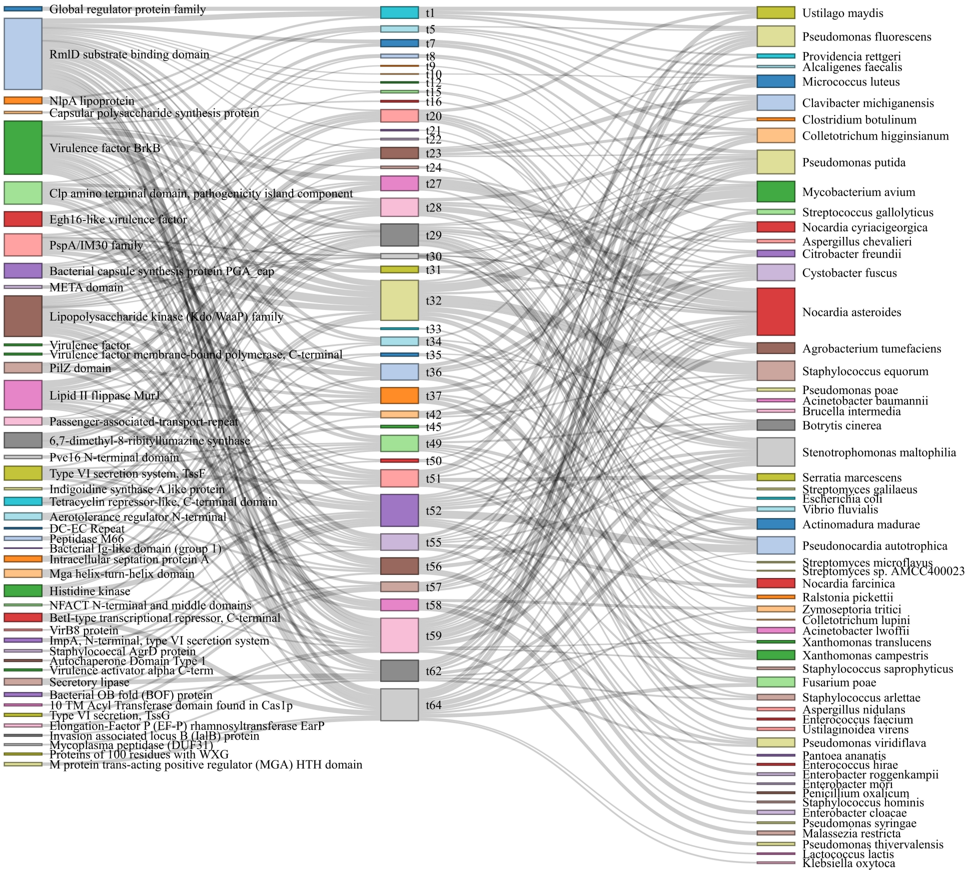


**Supporting Figure 22.** Host microorganisms carrying virulence traits with known pathogenicity to humans, animals, and plants found in air samples. The length of the rectangles indicates the RPKM values. From left to right, the rectangles represent virulence-related traits, sampling day (time series), and potential pathogens. For visual convenience, RPKM values were not depicted, but for reference, the total abundance of RmlD substrate binding domain (blue colored rectangle left-top) is 89.26 RPKM.

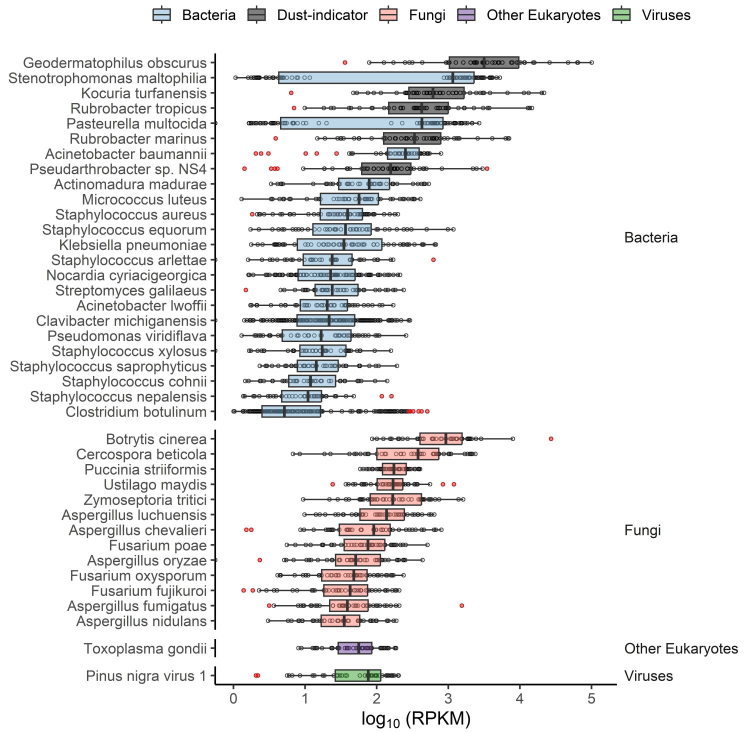


**Supporting Figure 23.** Abundance of contigs associated with known pathogenicity to humans, animals, and plants found in air samples. Boxplot center lines represent the median values, while the lower and upper hinges correspond to the first and third quartiles (25th and 75th percentiles). The lower and upper whiskers extend from the hinges to the lowest and largest values within 1.5 times the Interquartile Range (IQR) from the hinge. Outliers are displayed as red-colored dots.

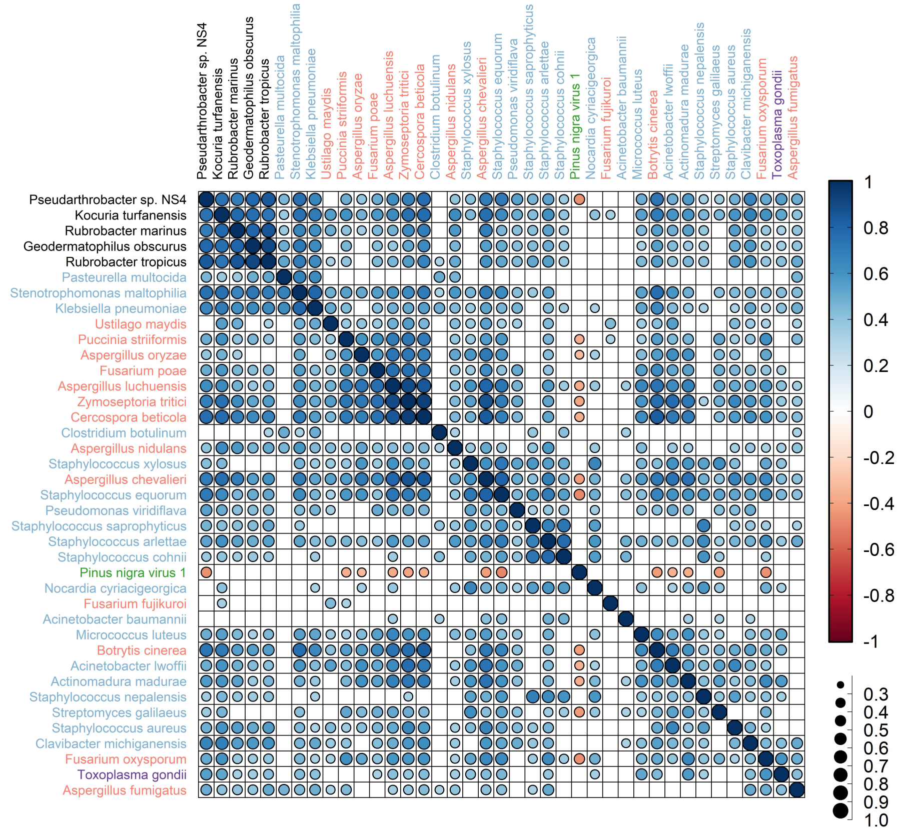


**Supporting Figure 24.** Correlation matrix of Spearman's coefficients showing co-occurrence patterns between potential pathogens and dust-indicator taxa. Correlations with an absolute Spearman's coefficient ($\rho$) value above 0.3 and a P-value below 0.05, adjusted by the Benjamini–Hochberg method, were considered statistically significant. The visual representation uses color-coded correlation coefficient values and different-sized bins for ease of interpretation. Taxa names are color-coded: black for dust-indicator taxa, light blue for bacteria, light red for fungi, green for viruses, and magenta for other eukaryotes. The x-axis and y-axis ordering were determined through hierarchical clustering using Ward's method with the ward.D option in the hclust package. This method minimizes within-cluster variance and combines clusters based on the smallest between-cluster distance.

**Supporting Table 1 (Separate file).** Metagenomic sequencing results of air samples before trimming and quality assessment.

**Supporting Table 2 (Separate file).** Metagenomic sequencing results of air samples after trimming and quality assessment.

**Supporting Table 3 (Separate file).** Meteorological data and estimated origin of air masses.

**Supporting Table 4.** Adonis test results for environmental variables.

| | Variable | Df | SumOfSqs | R2 | F-value | P-value |
|---|---|---|---|---|---|---|
| Taxonomic Features | $PM_{10}$ | 1 | 0.510 | 0.098 | 5.537 | <0.001 |
| | Air mass | 2 | 0.483 | 0.093 | 2.620 | <0.001 |
| | Relative humidity | 1 | 0.118 | 0.023 | 1.280 | 0.210 |
| | Temperature | 1 | 0.484 | 0.093 | 5.253 | <0.001 |
| | Residual | 39 | 3.592 | 0.693 | | |
| | Total | 44 | 5.186 | 1.000 | | |
| Functional Features | $PM_{10}$ | 1 | 0.017 | 0.088 | 4.341 | 0.004 |
| | Air mass | 2 | 0.011 | 0.058 | 1.421 | 0.114 |
| | Relative humidity | 1 | 0.003 | 0.016 | 0.775 | 0.582 |
| | Temperature | 1 | 0.009 | 0.046 | 2.261 | 0.036 |
| | Residual | 39 | 0.154 | 0.792 | | |
| | Total | 44 | 0.194 | 1.000 | | |

Df (degrees of freedom) signifies the count of independent values available to estimate another parameter or statistic. SumOfSqs (sum of squares) denotes the sum of squared deviations of values from their mean. $R^2$ represents the proportion of variance in the dependent variable predictable from the independent variable(s). For instance, an $R^2$ value of 0.10 indicates that 10% of the total variance is explained by the specified variable.

**Supporting Table 5 (Separate file).** Results of Kraken2 taxonomic classification with Bracken refinement. It presents the CPM normalized classification results.

**Supporting Table 6 (Separate file).** Genome coverage and depth information of potential pathogens per chromosome.

**Supporting Table 7 (Separate file).** Genome coverage information for potential pathogens, binned by coverage depth across the entire genome.

**Supporting Table 8 (Separate file).** Genome coverage depth information for each specific position across the entire genome of potential pathogens.

**Supporting Table 9 (Separate file).** Results of KrakenUniq analysis. It presents the CPM normalized estimation of distinct minimizers per taxon in the input sequencing data.

**Supporting Table 10 (Separate file).** Abundance of antimicrobial resistance-contigs, normalized to RPKM.

**Supporting Table 11 (Separate file).** Abundance of virulence-related-contigs, normalized to RPKM.

**Supporting Table 12 (Separate file).** List of potential pathogens obtained from publicly available databases.

**Supporting Table 13 (Separate file).** Kraken taxonomic classification results before Bracken refinement.

**Supporting Movie 1.** Time-averaged maps of dust column mass density (kg $m^{-2}$). The data covers the time period from 26 March (00:30) to 15 April 2022 (18:30). Each frame displays the date and hour at the center top. The color code indicates values, ranging from minimum (blue) to maximum (red).

**Supporting Movie 2.** Time-averaged maps of dust column mass density (kg $m^{-2}$). The data covers the time period from 16 April (00:30) to 30 April 2022 (18:30). Each frame displays the date and hour at the center top. The color code indicates values, ranging from minimum (blue) to maximum (red).

**Supporting Movie 3.** Time-averaged maps of dust column mass density (kg $m^{-2}$). The data covers the time period from 1 May (00:30) to 15 May 2022 (18:30). Each frame displays the date and hour at the center top. The color code indicates values, ranging from minimum (blue) to maximum (red).

**Supporting Movie 4.** Time-averaged maps of dust column mass density (kg $m^{-2}$). The data covers the time period from 16 May (00:30) to 31 May 2022 (18:30). Each frame displays the date and hour at the center top. The color code indicates values, ranging from minimum (blue) to maximum (red).

#### References


1 Gusareva, E. S. *et al.* Microbial communities in the tropical air ecosystem follow a precise diel cycle. *Proc Natl Acad Sci U S A* **116**, 23299-23308 (2019). https://doi.org:10.1073/pnas.1908493116

2 Erkorkmaz, B. A., Gat, D. & Rudich, Y. Aerial transport of bacteria by dust plumes in the Eastern Mediterranean revealed by complementary rRNA/rRNA-gene sequencing. *Communications Earth & Environment* **4**, 24 (2023). https://doi.org:10.1038/s43247-023-00679-8

3 Acker, J. G. & Leptoukh, G. Online analysis enhances use of NASA Earth science data. *Eos, Transactions American Geophysical Union* **88**, 14-17 (2007). https://doi.org:10.1029/2007eo020003

4 Cohen, M. D. *et al.* NOAA's HYSPLIT Atmospheric Transport and Dispersion Modeling System. *Bulletin of the American Meteorological Society* **96**, 2059-2077 (2015). https://doi.org:10.1175/bams-d-14-00110.1

5 Rolph, G., Stein, A. & Stunder, B. Real-time Environmental Applications and Display sYstem: READY. *Environmental Modelling & Software* **95**, 210-228 (2017). https://doi.org:10.1016/j.envsoft.2017.06.025

6 Wood, D. E., Lu, J. & Langmead, B. Improved metagenomic analysis with Kraken 2. *Genome Biol* **20**, 257 (2019). https://doi.org:10.1186/s13059-019-1891-0

7 Morgulis, A., Gertz, E. M., Schäffer, A. A. & Agarwala, R. A fast and symmetric DUST implementation to mask low-complexity DNA sequences. *J Comput Biol* **13**, 1028-1040 (2006). https://doi.org:10.1089/cmb.2006.13.1028

8 Lu, J., Breitwieser, F. P., Thielen, P. & Salzberg, S. L. Bracken: estimating species abundance in metagenomics data. *Peerj Comput Sci* **3**, e104 (2017). https://doi.org:10.7717/peerj-cs.104

9 Breitwieser, F. P., Baker, D. N. & Salzberg, S. L. KrakenUniq: confident and fast metagenomics classification using unique k-mer counts. *Genome Biol* **19**, 198 (2018). https://doi.org:10.1186/s13059-018-1568-0

10 Sarafian, R., Nissenbaum, D., Raveh-Rubin, S., Agrawal, V. & Rudich, Y. Deep multi-task learning for early warnings of dust events implemented for the Middle East. *npj Climate and Atmospheric Science* **6**, 23 (2023). https://doi.org:10.1038/s41612-023-00348-9
